# Single-cell, Spatially-Resolved TCR Profiling Links T Cell Phenotype and Clonality in Human Tumors

**DOI:** 10.1101/2025.08.06.668925

**Authors:** Kelli A. McCord, Emerald Kan, Sean Hyslop, Amanda Y. Xia, Colby J. Hofferek, James S. Lewis, Andreas Wieland, David J. Hernandez, Vlad C. Sandulache, William H. Hudson

## Abstract

Despite remarkable success in melanoma and other cancers, immune checkpoint blockade has shown limited efficacy in head and neck squamous cell carcinoma (HNSCC), with durable responses in only ∼10% of patients. This is paradoxical given the abundant infiltration of CD8⁺ T cells in HNSCC, suggesting that current T cell profiling methods fail to fully capture functional anti-tumor immunity and that key immunosuppressive mechanisms remain poorly understood. To uncover mechanisms regulating T cell function *in situ*, we developed a multimodal profiling framework integrating high-dimensional flow cytometry, single-cell RNA- and TCR-sequencing, and spatial transcriptomics, including spatially-resolved TCR profiling at single-cell resolution. Applying this approach to paired blood and tumor samples from 27 HNSCC patients enabled single-cell resolution mapping of T cell clonality, transcriptional state, and spatial localization within intact tumors. Tumors with similar clinical features and immune profiles by flow cytometry and scRNA-seq exhibited markedly different spatial immune architectures. Within the T cell compartment, stem- and memory-like T cells localized within immune-rich niches, while exhausted cells were broadly dispersed in tumor-rich regions. Targeted spatial TCR mapping revealed that tumor-enriched TCR clones were distributed throughout the tumor yet adopted distinct, location-dependent phenotypes, indicating that both antigen specificity and spatial cues shape T cell differentiation and function in human tumors. These findings reveal fundamental principles of T cell organization in solid tumors and establish a versatile platform for spatially-resolved, antigen-specific T cell profiling.

## Introduction

Head and neck squamous cell carcinoma (HNSCC) encompasses malignant tumors that develop in the sinonasal cavity, oral cavity, oropharynx, hypopharynx, and larynx^1^. Accounting for nearly 5% of cancer-related deaths^2, 3^, HNSCC represents a major global health concern. Standard-of-care therapy for HNSCC includes a combination of surgery, external beam radiotherapy and cytotoxic chemotherapy, which although effective in 50-75% of patients is associated with substantial morbidity^4–6^. Since 2016, PD-1 pathway blockade has been approved for use in recurrent/metastatic HNSCC^7, 8^ and, more recently, as a neoadjuvant therapy for resectable, locally advanced disease^9^. However, in both settings, only ∼10% of patients mount a durable response to PD-1 pathway blockade^8, 9^. This limited efficacy is striking given the presence of tumor-specific T cells within HNSCC tumors, particularly in human papillomavirus-associated (HPV^+^) disease^10^. This disconnect between tumor-specific T cell infiltration and therapeutic success highlights a fundamental gap in our knowledge of what drives effective anti-tumor immunity.

Paradoxically, while CD8^+^ T cells are the intended targets of current immune checkpoint blockade (ICB) therapies, we and others have found their presence to be associated with improved outcomes after standard-of-care therapy^11–13^, but not consistently with ICB response^14–17^. Instead, emerging evidence suggests that the spatial organization of immune components, particularly T cells, within the tumor microenvironment (TME) is a key factor in determining immunotherapy outcomes^10, 18–35^. However, many of these studies have lacked information on T cell specificity, an important consideration given the high infiltration of bystander (non-cancer-specific) T cells into human tumors^20, 36–39^. To overcome this limitation, additional studies have integrated T cell receptor (TCR)-sequencing with spatial transcriptomics to identify the location of antigen-specific T cells within the TME^19, 29, 40–42^. However, these current approaches lack the resolution to map individual T cell clones at single-cell resolution, limiting our understanding of how clonal identity, transcriptional state, and spatial positioning intersect to influence T cell function and differentiation in the TME.

To overcome current limitations in spatial T cell profiling and better resolve T cell biology within the TME, we applied a multimodal approach integrating high-dimensional flow cytometry, single-cell RNA and TCR sequencing, and spatial transcriptomics on paired tumor and blood samples from 27 HNSCC patients. Tumor-infiltrating lymphocytes (TILs) displayed distinct phenotypes and TCR repertoires compared to circulating T cells, highlighting that blood-based profiling alone fails to capture the complexity of the intratumoral immune response. Spatial transcriptomics revealed that tumors with similar clinical features and immune infiltration profiles by flow cytometry and scRNA-seq exhibited markedly different T cell spatial architectures, indicating that spatial organization provides orthogonal information not captured by conventional profiling. By integrating transcriptional phenotype with spatial location, we found that stem- and memory-like T cells localized within immune-rich niches, while exhausted cells were diffusely distributed throughout the tumor. Without clonal resolution, these spatial patterns could reflect differences in specificity - such as bystander versus tumor-reactive clones - rather than true phenotypic plasticity. To directly link clonal identity to spatial context, we mapped patient-specific TCRs using targeted spatial transcriptomics probes. T cells sharing the same TCR exhibited substantial transcriptional heterogeneity, spanning exhausted and more plastic (‘stem-like’) states, and were distributed across distinct tumor regions. Antigen specificity also shaped T cell distribution, with tumor-enriched clones exhibiting distinct neighboring cell types and gene expression profiles compared to bystander T cells. These findings demonstrate that T cell differentiation within the TME is not dictated by clonal identity alone, but by dynamic interplay between antigen specificity and environmental cues. Together, our approach enables antigen-specific, spatially-resolved T cell profiling at single-cell resolution and reveals new principles of T cell organization and differentiation in HNSCC tumors.

## Results

### Flow cytometry reveals compartment-specific and inter-patient immune heterogeneity in HNSCC

Over 14 months, we prospectively collected fresh tumor and blood samples from 27 patients with HNSCC (**Table S1, S2**). A portion of each tumor was embedded in OCT for spatial transcriptomics, while peripheral blood mononuclear cells (PBMCs) and TILs were isolated and viably cryopreserved (**Figure 1A**). Paired samples from all patients and PBMCs from five healthy donors were thawed, stained with a 31-color fluorescent antibody panel (**Table S3**), and analyzed on a spectral cell sorter in a single batch, enabling simultaneous high-dimensional immune profiling and isolation of viable T cells for downstream scRNA-seq and TCR-seq.

**Figure 1.**
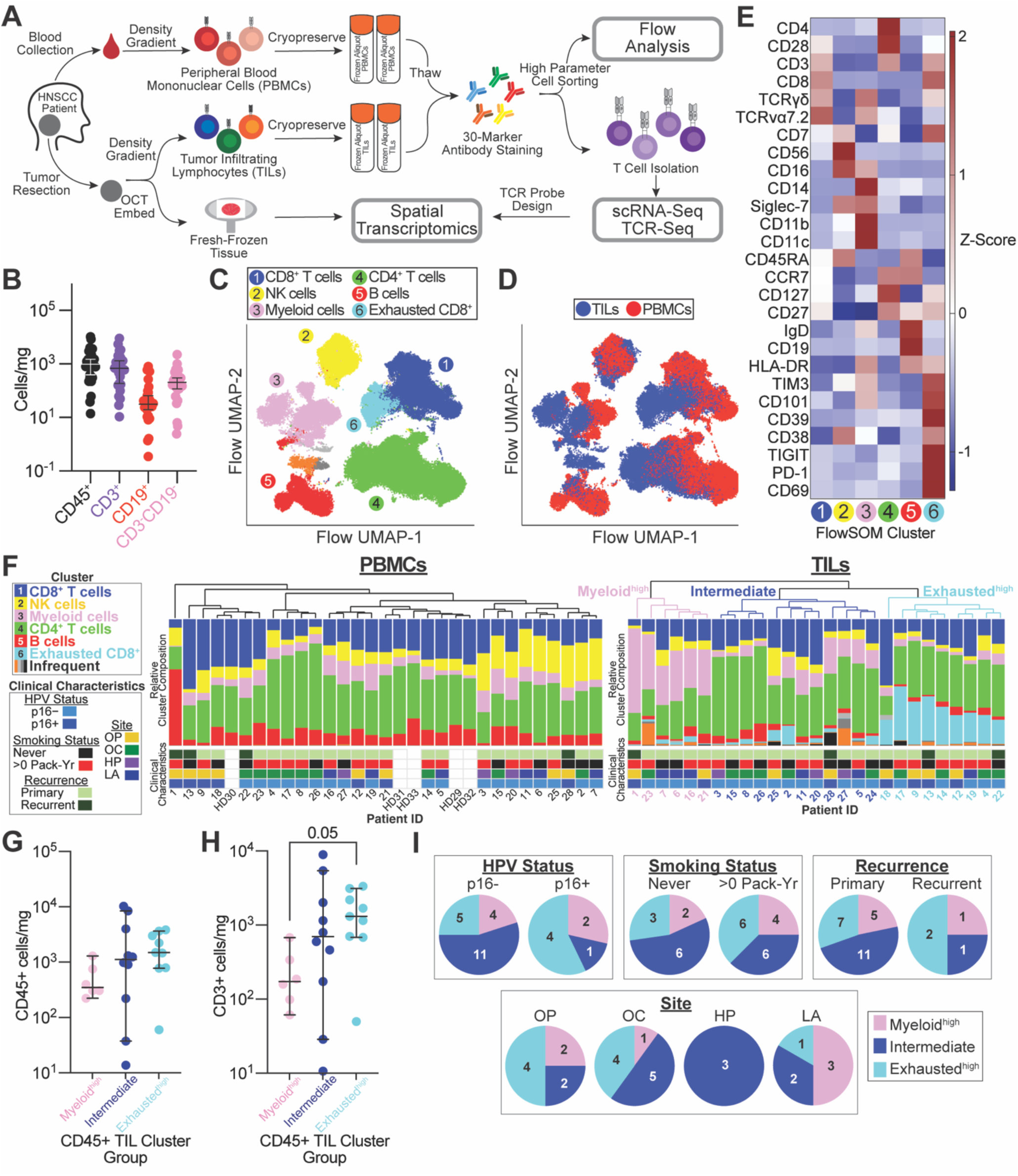
Distinct immune compositions exist between PBMCs and TILs across HNSCC patients. **(A)** Experimental workflow combining high-parameter flow cytometry, scRNA-seq, and spatial transcriptomics performed on paired blood and tumor samples collected from 27 patients with head and neck squamous cell carcinoma (HNSCC). **(B)** Quantification of CD45⁺ immune cell density in tumors, grouped by major immune subtypes. **(C)** UMAP of flow cytometry data from 27 patient PBMC and TIL samples, gated on live CD45⁺ cells and clustered using FlowSOM plugin on FlowJo. Cell populations comprising >1.1% of total CD45⁺ cells are annotated. **(D)** UMAP colored by tissue of origin (PBMCs vs. TILs). **(E)** Heatmap showing relative protein expression across FlowSOM-defined CD45⁺ cell clusters. **(F)** Relative composition of CD45^+^ FlowSOM-defined clusters per patient for PBMCs (left) and TILs (right), with healthy donor (HD) PBMCs included for reference. Patients were grouped by hierarchical clustering of TIL immune composition, which identified three categories: myeloid^high^, exhausted^high^, and intermediate. Clinical characteristics are annotated below each patient. Tumor sites include oropharynx (OP), oral cavity (OC), hypopharynx (HP), and larynx (LA). Smoking is measured in pack years (Pack-Yr). **(G, H)** CD45⁺ **(G)** and CD3⁺ cells **(H)** per milligram of tumor stratified by CD45^+^ flow groups. **(I**) Distribution of CD45⁺ flow groups within patients stratified by clinical characteristics. Error bars on graphs B, H, I are median with 95% confidence interval (CI). Statistical testing in **(G-H)** was performed using the Kruskal-Wallis test, corrected with multiple comparisons.

T cells were the predominant immune cell population in most tumors (**Figure 1B**), although immune composition varied among patients and did not correlate with clinical features such as p16 status, tumor site, smoking history, or recurrence (**Figure S1A-D**). Unsupervised clustering of CD45^+^ cells identified nine immune cell populations (**Figure 1C, S1F**), with clear differences in abundance between PBMC- and TIL-derived immune subsets (**Figure 1D, S1E**). We annotated the six most abundant clusters (>1.1%) based on differential protein expression (**Figure 1E, S1G**) and comparison to conventional flow cytometry gating strategies (**Figure S1H**). NK cells and B cells were enriched in the blood while TILs contained higher proportions of myeloid and exhausted CD8^+^ T cells (**Figure S1E**). This pattern aligns with prior studies demonstrating that the TME supports accumulation of dysfunctional or immunosuppressive immune cells, while the circulating immune compartment is dominated by less immunosuppressive and antigen-experienced subsets^43–47^.

To quantify interpatient heterogeneity in immune composition, we performed hierarchical clustering of patients based on the relative abundance of CD45⁺ subsets within their TILs (**Figure 1F**). This analysis identified three distinct immune profiles, hereafter referred to as CD45^+^ flow groups: a myeloid^high^ group enriched in myeloid cells, an exhausted^high^ group enriched in exhausted CD8⁺ T cells, and an intermediate group with lower frequencies of both myeloid cells and exhausted CD8⁺ T cells, and a relative increase in CD4⁺ T cells and/or non-exhausted CD8⁺ T cells. Tumors in the myeloid^high^ group contained lower absolute numbers of CD45^+^ and CD3^+^ immune cells compared to the exhausted^high^ group (**Figure 1G-H**). Clinically, exhausted^high^ tumors were more common among p16^+^ cases, smokers, recurrent tumors, and tumors resected from the oropharynx or oral cavity (**Figure 1I**). These associations support a connection between immune landscapes and specific clinical features, while also demonstrating that immune heterogeneity in HNSCC is shaped by factors beyond conventional clinical classifications.

To further dissect heterogeneity within the T cell compartment, we performed unsupervised clustering of circulating and tumor-infiltrating CD4⁺ and CD8⁺ T cells (**Figure S2, S3**). Naïve T cells were enriched in PBMCs, while TILs contained higher proportions of regulatory CD4⁺ T cells, exhausted CD8⁺ T cells, and CD8^+^ memory-like cells, again highlighting clear differences between circulating and tumor-infiltrating cells. CD4^+^ T_regs_ and exhausted CD8⁺ T cells were most abundant in the exhausted^high^ group, reinforcing the distinct immune landscape that defines each CD45⁺ flow group.

### scRNA-seq defines transcriptionally and clonally distinct tumor-infiltrating T cell states

To resolve the transcriptional complexity underlying the phenotypic differences observed by flow cytometry, we performed scRNA-seq on tumor-infiltrating T cells isolated during high-dimensional flow profiling. Nearest neighbor clustering of 251,708 high-quality single T cells revealed 14 transcriptionally distinct populations (**Figure 2A-B**). Clusters 1-3 (1.2% of sequenced cells) were excluded from downstream analysis due to signatures indicative of doublets and/or dying cells. The remaining 11 clusters exhibited gene expression patterns of well-defined T cell subsets, such as *CCR7*^+^ naïve/central memory cells (N/Tcm; cluster 10), *FOXP3*^+^ regulatory cells (T_regs_; cluster 12), and *PDCD1*^+^ (PD-1^+^) *ENTPD1^+^* (CD39^+^) exhausted cells (clusters 13 and 14) (**Figure 2C**).

**Figure 2.**
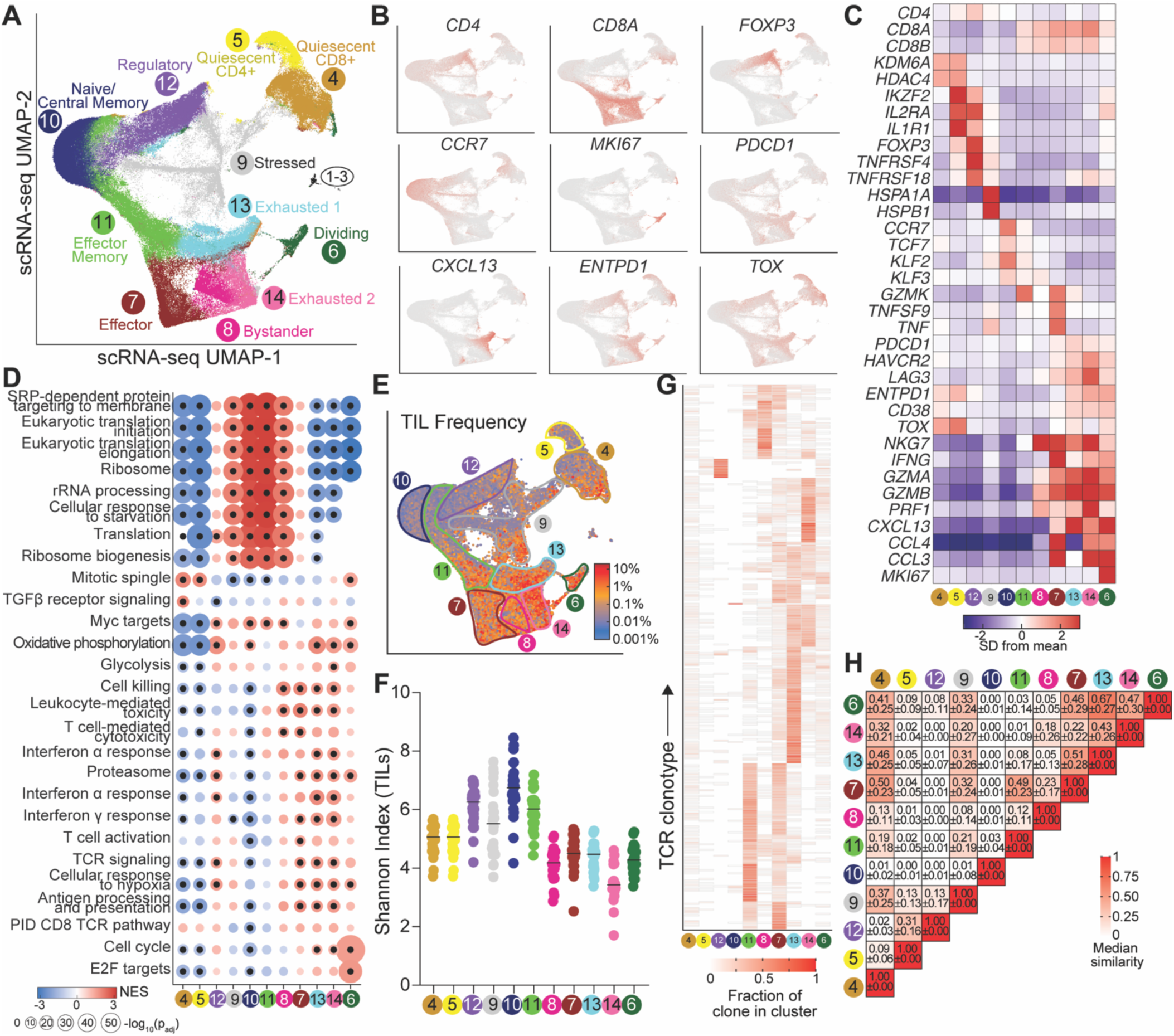
scRNA-seq of tumor-infiltrating T cells reveals transcriptionally diverse T cell states in HNSCC. **(A)** Uniform manifold approximation and projection (UMAP) of scRNA-seq data from tumor-infiltrating T cells, annotated by nearest-neighbor clustering. **(B)** Expression of canonical T cell subset markers. **(C)** Average expression of selected T cell markers across scRNA-seq clusters. **(D)** Gene set enrichment analysis (GSEA) based on differential gene expression in each cluster versus all others. Pathways are drawn from Reactome (RE), Kyoto Encyclopedia of Genes and Genomes (KEGG), Hallmark (HK), and Gene Ontology (GO) gene sets. **(E)** Paired single-cell TCR sequencing of tumor-infiltrating T cells. Each cell is colored by the frequency of its TCR within the patient tumor. **(F)** Shannon diversity index of TCRs within each cluster. Each point represents a single patient. **(G)** Distribution of high-frequency TCR clones across clusters. Each row represents a TCR clone present at >0.5% frequency within a patient; columns indicate the proportion of that clone found in each cluster. **(H)** Quantification of TCR overlap between scRNA-seq phenotypic clusters as measured by the Morisita-Horn similarity index (median ± SD).

Exhausted clusters were enriched for genes involved in metabolism, proliferation, cytotoxicity, TCR signaling, and hypoxia response, while displaying reduced expression of ribosomal and protein trafficking pathways. Quiescent clusters lacked activation signatures but showed enrichment for TGFβ signaling, consistent with a suppressed phenotype. These transcriptional differences underscore the diversity of T cells in the TME and raise the possibility that differentiation status may play key roles in shaping their functional states and spatial organization (**Figure 2D**).

To investigate clonal dynamics between these subsets within the TME, we performed paired single-cell TCR-seq, capturing both the α/β and the less commonly profiled γ/δ TCR chains. Exhausted clones represented the most expanded T cell populations and showed the lowest clonal diversity across all subsets (**Figure 2E-F**). N/T_cm_ cells were the most diverse, while T_regs_ and effector memory (T_em_) cells showed intermediate levels of clonal expansion. Dividing cells also showed low TCR diversity, indicating selective entry of tumor-infiltrating clones into the cell cycle. TCR overlap analysis revealed the greatest clonal sharing between the two exhausted clusters and between T_em_ and effector clusters (**Figure 2G-H**). Both the T_reg_ cluster and bystander cluster, which we later characterize as enriched for non-tumor-specific CD8^+^ T cells, displayed minimal TCR overlap with other clusters, suggesting they are distinct from the broader differentiation trajectories of intratumoral T cells.

### Circulating and tumor-infiltrating T cells exhibit divergent clonal and phenotypic landscapes

To understand compartment-specific differences in T cell clonal architecture, we performed α/β TCR-sequencing of CD4^+^ and CD8^+^ T cells sorted from 27 HNSCC patient-matched PBMCs and five healthy donors. CD8⁺ T cells were more clonally expanded than CD4⁺ T cells in both patient and healthy donor PBMCs, with the lowest diversity observed in HNSCC patient-derived CD8⁺ T cells (**Figure 3A**). This reduced diversity in CD8^+^ T cells extended to TILs (**Figure 3B**) and was independent of p16 status and smoking history (**Figure S4A-B**). Overall, TILs were significantly more clonally expanded than matched PBMCs (**Figure 3C**). Clonal overlap between PBMCs and TILs was limited but higher in CD8^+^ than CD4^+^ T cells (median Morisita-Horn similarity index, 0.18 vs 0.08; **Figure 3D**), with no significant differences by p16 or smoking status (**Figure S4C-D**). When stratified by CD45⁺ flow group, we observed that exhaustedʰⁱᵍʰ tumors had reduced clonal overlap between PBMCs and TILs (**Figure 3E**), indicating that high intratumoral exhaustion is associated with greater divergence from circulating TCR repertoires.

**Figure 3.**
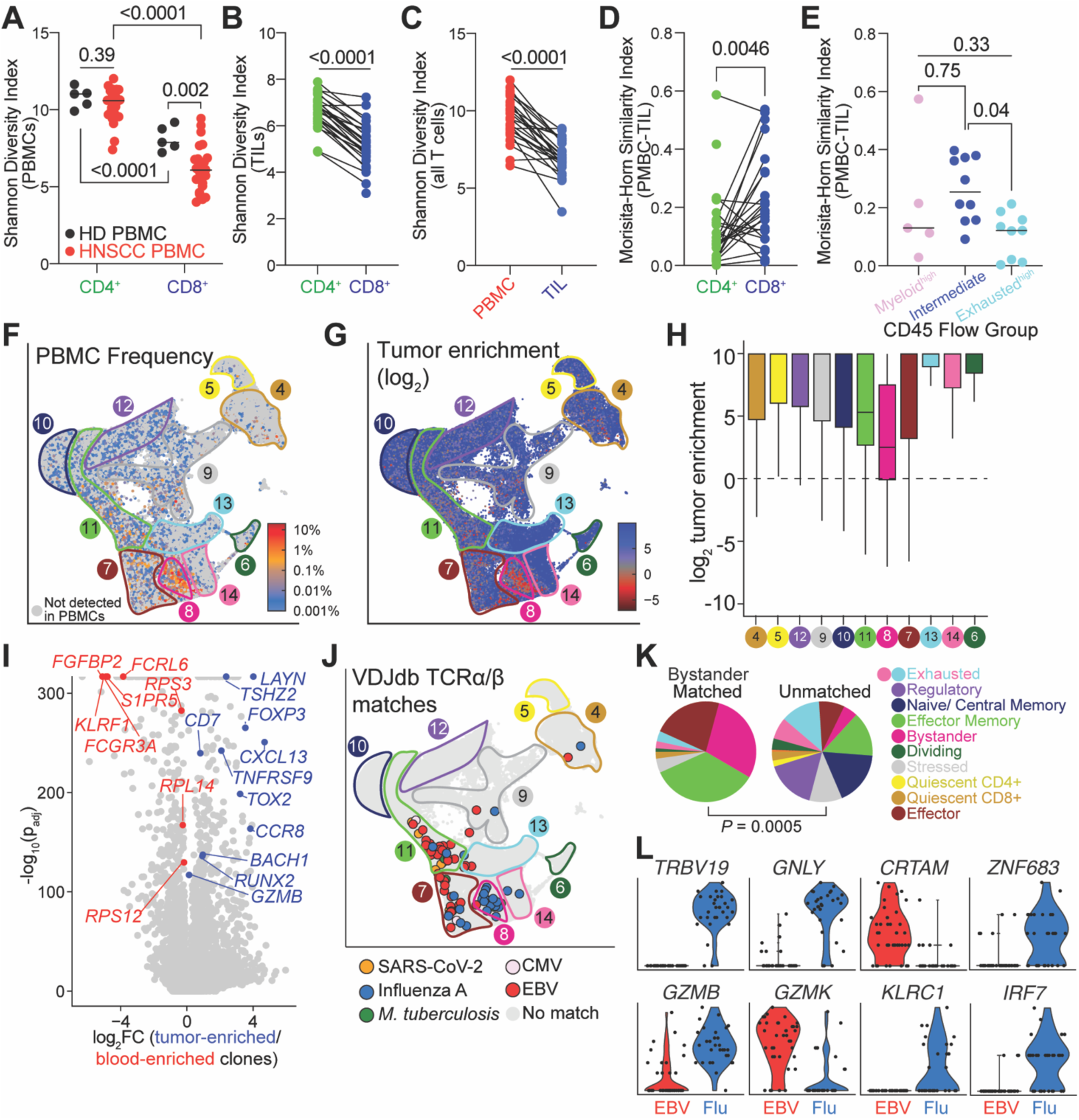
TCR-sequencing identifies distinct phenotypes of tumor-enriched and bystander TCRs within the HNSCC TME. **(A)** TCR diversity of CD4⁺ and CD8⁺ T cells isolated from healthy donor PBMCs (HD, n =5) and HNSCC patient PBMCs (PBMC, n=26). Clonal diversity was quantified using the Shannon diversity index. **(B)** Shannon diversity index of TCRs from CD4⁺ and CD8⁺ TIL TCRs within each patient (n=27). **(C)** Shannon diversity index of all TCRs found in patient-matched PBMCs (n=26) and TILs (n=27). **(D)** Morisita-Horn similarity index values comparing TCR overlap between HNSCC patient PBMCs and TILs for CD4^+^ T cells and CD8^+^ T cells. **(E)** Morisita-Horn values comparing overall TCR overlap between PBMCs and TILs, stratified by patient CD45^+^ flow groups. **(F)** scRNA-seq UMAP of tumor-infiltrating T cells, colored by the frequency of each cell’s TCR in matched peripheral blood. Clones not detected in PBMCs are shown in gray. **(G)** scRNA-seq UMAP of tumor-infiltrating T cells, colored by the tumor enrichment of each cell’s TCR (ratio of TCR frequency in TILs to PBMCs). **(H)** Tumor enrichment of TCR clones by subcluster. Boxplots show log₂ tumor enrichment of clones across subclusters; center line = median, box = interquartile range (IQR), whiskers = ±1.5×IQR. Horizontal dashed line indicates no enrichment (0). **(I)** Differential gene expression between tumor-enriched and blood-enriched clones; positive log_2_FC values indicate genes more highly expressed in tumor-enriched clones. **(J-K)** Phenotype of cells with VDJdb-annotated TCRs specific for SARS-CoV-2, CMV, Influenza A, EBV, and *M. tuberculosis* (colored), or unmatched TCRs (gray). **(L)** Expression of selected genes within cells containing TCRs specific for EBV (red) or influenza (red). Statistical tests: **(A)** mixed-effects analysis with patient-matched pairing and Fisher’s LSD test; **(B-D)** paired t-test; **(E)** one-way ANOVA with Tukey’s post hoc correction; **(K)** Fisher’s exact test.

Within tumor-infiltrating T cells, TCRs abundant in circulation were predominantly found in cluster 8, which we thus termed the ‘bystander’ cluster (**Figure 3F**). Since gene expression clustering was performed in a TCR-agnostic manner, the highly selective enrichment of circulating TCRs in this phenotypically-defined cluster is particularly striking. Tumor enrichment - defined as the ratio of each TCR’s frequency in TILs to its frequency in PBMCs - was lowest in this cluster (**Figure 3G-H**). T cells with tumor-enriched clones upregulated regulatory, effector, and exhaustion-associated genes (e.g., *FOXP3*, *GZMB*, *CXCL13)*, while blood-enriched clones showed increased expression of ribosomal (*RPS3*, *RPL14*), migration (*S1PR5*), and NK-like (*FCGR3A*) transcripts (**Figure 3I**). Notably, *FGFBP2* expression was a strong marker of tumor-infiltrating T cells with blood-enriched clonotypes. Given that its expression has been previously linked with ICB response in lung cancer^48^, FGFBP2 may serve as a surrogate for T cell infiltration capacity into the TME.

To investigate the antigen specificity of TIL-derived TCRs, we cross-referenced tumor-infiltrating clonotypes with the VDJdb database^49^. TCRs matching known microbial antigen reactivities were primarily found within the bystander, effector, and T_em_ clusters (**Figure 3J-K**). Interestingly, TCRs specific for EBV and influenza A mapped to distinct clusters and exhibited divergent gene expression programs (**Figure 3J,L**), suggesting that pathogen-specific immune imprinting may contribute to heterogeneity of bystander T cells across patients and, in turn, influence the overall immune landscape of the TME.

Given the marked heterogeneity of T cell phenotypes within tumors, we extended our analysis to include unconventional T cell subsets, γδ and MAIT cells, to understand their contribution to the immune landscape in HNSCC. γδ T cells predominantly mapped to exhausted, memory-like, and bystander scRNA-seq clusters, exhibited an effector/NK-like transcriptional profile marked by expression of *NCAM1, GZMB*, and KLR family members, and were more abundant in circulation than in tumors (**Figure S4E-K**). In addition to reduced transcript expression of *CD8A* and *CD4*, flow cytometry confirmed that many tumor-infiltrating γδ T cells were CD4⁻CD8⁻ (**Figure S4I-J**). However, despite lacking CD8 expression, γδ T cells primarily localized to CD8⁺-enriched clusters, reflecting a cytotoxic phenotype. In contrast, MAIT cells were broadly distributed across all T cell clusters and expressed canonical markers^50–52^ (e.g. *KLRB1*, encoding CD161) while downregulating stemness-associated genes^53^ (e.g. *SELL*, encoding CD62L) (**Figure S4L-N**). Compared to non-MAIT cells, MAIT cells showed lower tumor enrichment by TCR-seq, potentially reflecting their specificity for microbial rather than tumor antigens^50^. Flow cytometry confirmed that circulating MAIT cells were significantly depleted in HNSCC patients relative to healthy donors (**Figure S4O-Q**), consistent with prior findings in other cancers^54–57^.

### Spatial transcriptomics defines the cellular landscape of HNSCC at single-cell resolution

While flow cytometry and scRNA-seq revealed striking heterogeneity in the phenotype and clonotypes of tumor-infiltrating T cells in HNSCC, they do not provide information about how these populations are spatially organized within the tumor. To address this, we performed Xenium spatial transcriptomics on tumor sections from ten of the patients previously profiled by flow cytometry and scRNA-seq, including seven patients processed in technical duplicates. Hematoxylin and eosin (H&E) staining was performed to determine tissue architecture (**Figure 4A**); nuclear, boundary, and interior RNA imaging enabled accurate cell segmentation within the complex HNSCC TME (**Figure 4B-C**).

**Figure 4.**
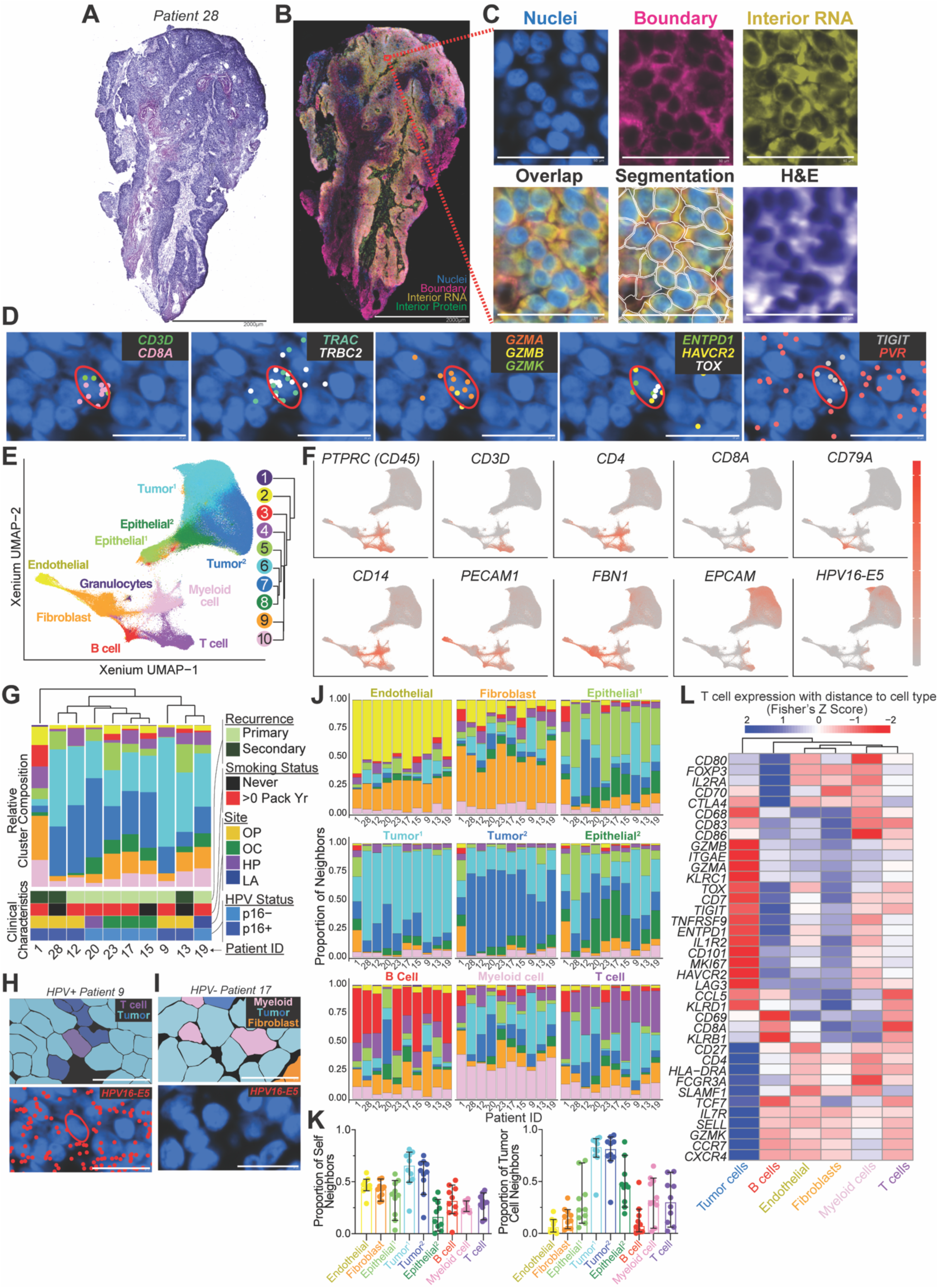
Spatial transcriptomic profiling of the HNSCC TME permits single-cell resolution of gene expression and identifies location-dependent T cell gene expression profiles. **(A-C)** Representative H&E staining (**A**), fluorescent staining for cell segmentation of nuclei (blue), membrane boundaries (pink), and cytoplasmic RNA (yellow) (**B**), and software-based cell segmentation (C) of a HNSCC tumor section. Scale bar represents 50 µm. **(D)** Spatial localization of selected transcripts within a representative tissue section from patient 9, shown with a 20 µm scale bar. **(E)** UMAP and nearest neighbor clustering of Xenium spatial transcriptomic data from 17 tumor sections across 10 patients. **(F)** Expression of selected canonical genes projected onto the Xenium UMAP. **(G)** Relative abundance of cells in each Xenium cluster across patients with annotated clinical characteristics. **(H-I)** Spatial mapping of Xenium-defined cell clusters projected onto tissue images. Cells are colored by transcriptional identity as defined by clustering of Xenium gene expression data. HPV16-E5 oncoprotein transcript expression is shown in a representative HPV⁺ (**H**) and HPV⁻ (**I**) tumor. Scale bar represents 20 µm. **(J)** Proportion of neighboring cell types, defined as the five nearest cells within a 15µm radius, for each cell population across patients. **(K)** Proportion of each cell type’s neighbors that are the same cell type (left) or tumor cells (right). Each dot represents one patient and error bars indicate median with 95% CI. **(L)** Correlation between T cell gene expression and their proximity to the indicated cell type.

Across all 17 sections, we measured the expression of 477 transcripts at single-cell resolution. This enabled precise cell-type identification, exemplified by a representative CD8⁺ T cell co-expressing *CD3D* and *CD8A*, confirming T cell identity, and *TRAC* and *TRBC2*, indicating the presence of an αβ TCR (**Figure 4D**). Effector potential was marked by expression of *GZMA* and *GZMB*, while *ENTPD1*, *HAVCR2*, and *TOX* identified an exhausted phenotype. Expression of the inhibitory receptor *TIGIT* alongside neighboring cells expressing its ligand, *PVR*, revealed putative immunoregulatory interactions. This resolution further enabled classification of T cell subsets, including *FOXP3⁺ IL2RA⁺* regulatory T cells and memory or stem-like populations marked by *TCF7*, *CCR7*, and *IL7R* (**Figure S5A-D**).

Using gene expression profiles measured by Xenium spatial transcriptomics from all 17 samples, we performed UMAP projection and nearest-neighbor clustering, identifying ten distinct gene expression clusters readily annotated as T cells, B cells, myeloid cells, endothelial cells, fibroblasts, granulocytes, and four epithelial subsets (two malignant/tumor, two normal; **Figure 4E-F**, **Figure S6A**). All clusters were present in every patient, but relative proportions varied, highlighting inter-patient heterogeneity of TME contents (**Figure 4G**). Cells from the tumor clusters were the predominant population in nine of ten patients. Patient 1, who had concurrent chronic lymphocytic leukemia, showed an atypical profile characterized by a higher proportion of B cells and minimal epithelial content. Cluster composition did not correlate with other clinical factors (**Figure 4G**). In addition to pathologist review of H&E staining, when possible we identified tumor cells using custom Xenium probes targeting *HPV16* gene products, which were selectively expressed in *HPV⁺* tumors (**Figure 5H-I**). Other cell type assignments were validated by overlaying gene expression clusters on tissue images and confirming localization of lineage-defining transcripts (**Figure S6B**). Lineage-specific markers remained relatively confined to the appropriate cells, supporting the accuracy of cell type classification and cell segmentation in the complex HNSCC TME.

**Figure 5.**
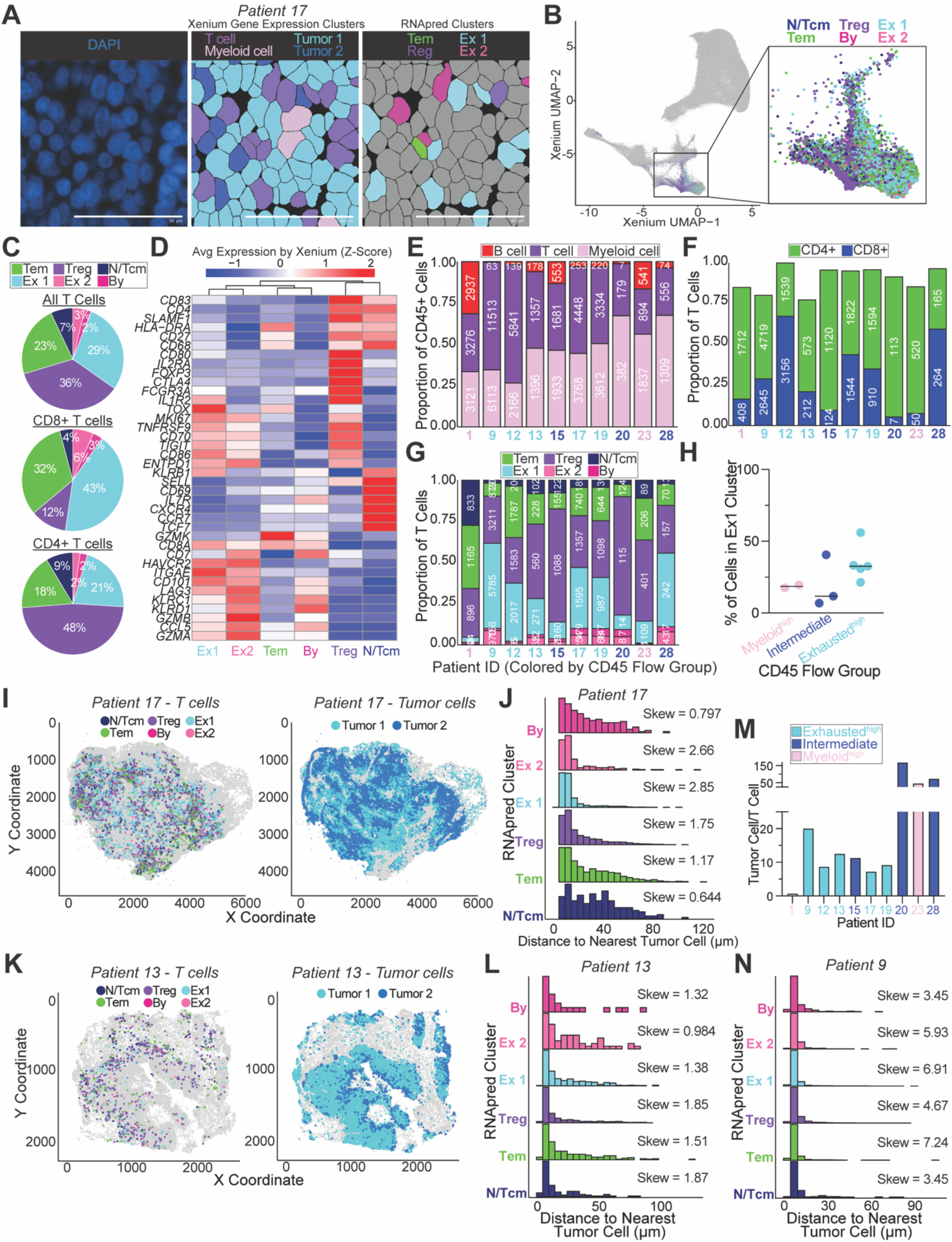
Integration of scRNA-seq and spatial transcriptomics data enables high-resolution phenotypic and spatial classification of T cells. **(A)** Representative tumor section, with nuclei visualized by DAPI (left) and segmented cells overlaid by either Xenium gene expression clusters (middle) or scRNA-seq-derived RNA_pred_ T cell phenotypes (right). Scale bar represents 50 µm. **(B)** Xenium UMAP colored by RNA_pred_ cluster assignments. **(C)** Distribution of RNA_pred_ clusters among all (top), CD8⁺ (middle), and CD4⁺ (bottom) T cells present within integrated Seurat object of 20,000 cells per patient. **(D)** Relative (Xenium) gene expression of selected T cell genes across RNA_pred_-defined clusters. (**E-G**) Relative roportion of CD45⁺ cells **(E)**, CD4⁺/CD8⁺ T cells **(F)**, and RNA_pred_ T cells **(G)** per patient. Patient IDs are color-coded by CD45 flow group; absolute counts are labeled. Cells lacking or co-expressing *CD4* and *CD8A* transcripts were not subtyped. **(H)** Percentage of cells present within the RNA_pred_ Ex1 cluster stratified by CD45^+^ flow group. **(I)** Spatial distribution of T cells in patient 17 tumor sample colored by RNA_pred_ cluster (left) and spatial distribution of tumor cells (right). **(J)** Distribution of distances from every T cell to the nearest tumor cell **(J)** within patient 17, grouped by RNA_pred_ cluster. **(K)** Spatial positioning of T cells colored by RNA_pred_ cluster (left) and spatial distribution of tumor cells (right) within the tumor sample from patient 13. **(L)** Distribution of distances from every T cell to the nearest tumor cell **(J)** within patient 13, grouped by RNA_pred_ cluster. **(M)** Tumor-to-T cell ratio from Xenium data across patients, colored by each patient’s CD45⁺ flow cytometry group. **(N)** Distribution of distances to the nearest tumor cell for each T cell arranged by RNA_pred_ cluster in patient 9, with skewness values reported per cluster.

### Cell types exhibit distinct spatial interactions within the TME

To investigate spatial relationships in the TME, we analyzed the neighboring cell types of each major population (**Figure 4J-K**). Endothelial and tumor cells were most often adjacent to their own cell type, forming vascular and epithelial zones. Fibroblasts neighbored nearly all cell types as well as each other, forming fibroblast-rich zones that extend broadly across the tumor. T cells and myeloid cells showed greater proximity to tumor cells than other immune or stromal populations, suggesting preferential positioning within malignant regions rather than peripheral or vascular zones. B cells primarily neighbored each other, with limited contact with tumor cells. These patterns indicate distinct spatial niches for endothelial and epithelial cells, widespread fibroblast distribution, and tumor-associated positioning of T cells and myeloid cells.

To probe cell-cell communication, we applied CellChat analysis^58^ and found the strongest predicted interactions between T cells and myeloid cells, highlighting their central role in TME communication (**Figure S6C-D**). Key pathways included *PVR*-*TIGIT* signals between tumor and T cells, and *CD86*-mediated signaling from myeloid to T cells, revealing immunosuppressive and costimulatory axes operating within the tumor^59–62^. Given the limited gene panel used in Xenium analysis, these findings reflect a fraction of the full signaling repertoire between T cells and myeloid cells.

Building on these spatial and signaling relationships, we next asked whether T cell transcriptional states reflect their proximity to specific cell types. Correlations between T cell gene expression and their distance from specific cell types (**Figure 4L**) revealed that exhaustion-associated genes (*ENTPD1*, *HAVCR2*, *LAG3*, *PDCD1*), as well as effector (*GZMA*, *GZMB*) and activation markers (*HLA-DRA*, *SLAMF1*), were elevated in T cells positioned near tumor and myeloid cells, suggesting activation upon tumor antigen encounter. In contrast, naïve and memory-associated transcripts (*IL7R*, *SELL*, *CCR7, GZMK*) were enriched in T cells positioned farther from tumor and myeloid cells and closer to B cells, endothelial cells, and other T cells. Regulatory markers (*FOXP3*, *IL2RA*) were most elevated in T cells near fibroblasts, endothelial, and myeloid cells. Thus, T cells occupy diverse spatial contexts within the HNSCC TME - showing activation and exhaustion signatures near tumor and myeloid cells, regulatory features near fibroblasts, endothelium, and myeloid cells, and more naïve or memory-like programs in regions enriched for B cells, endothelial cells, fibroblasts, and other T cells.

### Integration of scRNA-seq with Xenium refines phenotypic classification of spatially resolved T cells

While providing rich spatial context, the limited transcript coverage of the Xenium assay constrains the classification of precise T cell subsets. To address this, we leveraged our patient-matched scRNA-seq T cell atlas to guide phenotypic inference from spatial transcriptomics data. Specifically, we trained a random forest classifier to predict scRNA-seq cluster identities based on the expression of genes shared between the scRNA-seq and Xenium data. This model assigned a predicted RNA-seq cluster (RNA_pred_) to each Xenium T cell, excluding clusters representing temporary T cell states (dividing and quiescent).

Because true cluster identities are not known for Xenium T cells, we assessed model accuracy using a held-out portion of the scRNA-seq data that was not used for model training and observed an overall accuracy of 75.6% (**Figure S7**). The RNA_pred_ identities were next projected onto tissue sections (**Figure 5A**) to visualize transcriptional states *in situ* and onto the Xenium UMAP (**Figure 5B**), where T_regs_ and exhausted subsets localized to distinct regions of the T cell cluster, indicating that this model preserved key transcriptional distinctions despite limited gene coverage.

Across samples, exhausted 1 (Ex1), T_em_, and T_reg_ RNA_pred_ subsets constituted most Xenium-detected T cells, followed by N/T_cm_, exhausted 2 (Ex2), and bystander populations (**Figure 5C**). CD8⁺ T cells detected by Xenium were predominantly Ex1 and T_em_, while nearly half of CD4⁺ T cells were T_regs_ (**Figure 5C**). Xenium gene expression profiling of RNA_pred_ clusters recapitulated key transcriptional features observed in scRNA-seq (**Figure 5D**). For example, Ex1 and Ex2 clusters expressed high levels of exhaustion markers including *GZMB* and *TOX*, T_em_ cells exhibited elevated *GZMK*, T_regs_ expressed *FOXP3* and *IL1R2*, and N/T_cm_ cells expressed *CCR7*, *IL7R*, and *SELL*.

Within each patient, we quantified overall immune cell composition (**Figure 5E**), the proportion of CD4⁺ and CD8⁺ T cells (**Figure 5F**), and the distribution of RNA_pred_-assigned T cell phenotypes from the Xenium data (**Figure 5G**). When stratified by CD45^+^ flow groups, the myeloid^high^ group, which had fewer exhausted CD8⁺ T cells by flow, also showed reduced CD8⁺ T cell content and lower exhausted proportions in Xenium (**Figure 5H**), reinforcing cross-platform consistency. Spatial projection of RNA_pred_ cells in patient 17 revealed distinct phenotypic organization and broad T cell distribution throughout the tumor, consistent with widespread infiltration (**Figure 5I**). Ex1 and Ex2 T cells within this patient were preferentially located near tumor cells whereas N/T_cm_, T_em_, and bystander T cells were more often positioned at a distance (**Figure 5J**). These findings align with our earlier gene expression correlation analysis (**Figure 4L**), where exhaustion-associated genes were elevated in T cells close to tumor cells. However, the overall spatial architecture of the TME influenced the observed spatial relationships between T cell phenotypes; for example, an immune excluded tumor (patient 13) did not follow this correlation pattern (**Figure 5K-L**). Similarly, in patients with a high tumor-to-T cell ratio, spatial relationships were less apparent, as nearly all T cells were inherently close to tumor cells, limiting the dynamic range of this analysis (**Figure 5M-N**). These findings highlight the high degree of spatial heterogeneity among samples that may appear otherwise similar by flow cytometry and/or scRNA-seq, emphasizing that spatial context can reveal biologically meaningful variation not apparent through conventional single-cell analyses.

### Spatial clustering identifies organized immune niches with distinct T cell phenotypes

Organized immune aggregates have been observed across various tumor types, including HNSCC, and are thought to coordinate antitumor responses through structured immune cell interactions^29, 63, 64^. To further dissect whether the phenotypically distinct T cells found within the TME form discrete structures, we systematically quantified immune clustering and assessed T cell composition within these regions.

Using Moran’s I, a statistic that quantifies spatial autocorrelation^65, 66^, we found significant clustering of immune cells compared to randomly-selected cells (**Figure 6A**). Within T cells, CD4⁺ cells, but not CD8^+^, showed increased spatial autocorrelation compared to random distributions (**Figure 6B**). Among RNA_pred_-defined T cell subsets, T_reg_ and T_em_ cells displayed the strongest self-clustering (**Figure 6C**). Despite each patient having varied T cell numbers and phenotypic heterogeneity, these trends were relatively conserved across all tissue sections.

**Figure 6.**
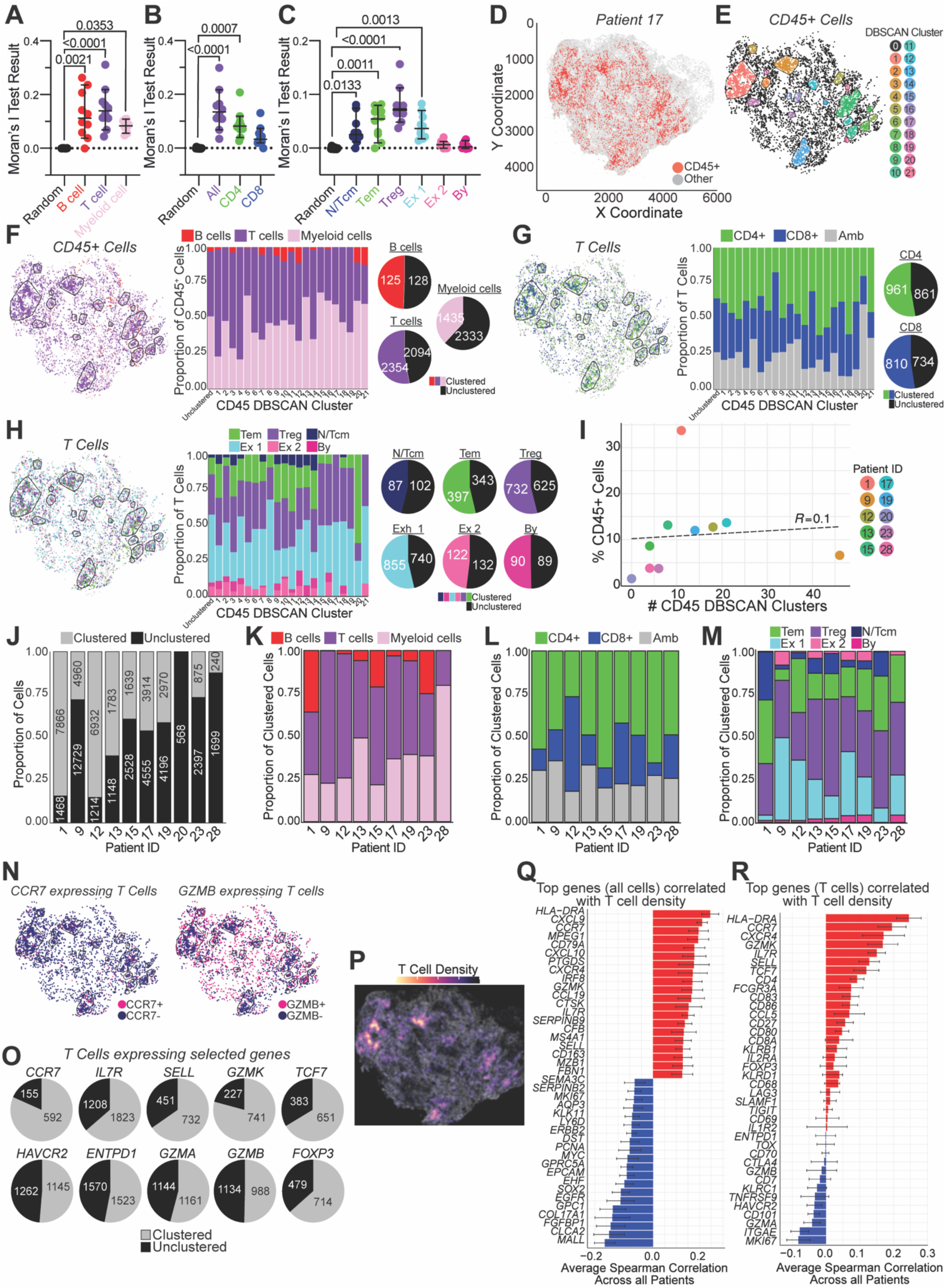
Spatial organization of immune cells is shaped by transcriptional phenotype and local gene expression. **(A-C)** Moran’s I spatial autocorrelation values for major immune lineages (T cells, B cells, myeloid cells) **(A)**, CD4⁺ and CD8⁺ T cells **(B)**, and RNA_pred_ T cell clusters **(C)**. Each point represents one patient. P values were measured using the Kruskal-Wallis test with Dunn’s multiple comparisons against a group of randomly-selected control cells. **(D)** Spatial distribution of CD45⁺ cells within a representative tissue section from patient 17. **(E)** DBSCAN spatial clustering of CD45⁺ cells in patient 17, using a minimum of 35 cells within an 80 µm radius. Unclustered CD45⁺ cells are shown in black. **(F-H)** Composition of DBSCAN clusters, grouped by broad immune cell classification (**F**), CD4^+^ and CD8^+^ T cells (**G**), and T cell RNA_pred_ phenotype (**H**). Left: Cell types with DBSCAN clusters overlaid on tissue. Middle: subset proportions within each cluster. Right: pie charts show fraction of each subset inside versus outside DBSCAN clusters. Ambiguous T cells (Amb) lack or co-express *CD4* and *CD8A* transcripts. **(I)** Number of DBSCAN-defined CD45⁺ clusters (x-axis) and the ratio of CD45⁺ cells to total cells per tissue section (y-axis) across patients, showing that immune cell abundance does not directly correlate with the number of spatial clusters. **(J)** Proportion of CD45⁺ cells found within versus outside of DBSCAN clusters across patients, highlighting interpatient variability in immune cell organization. **(K)** Immune cell composition of CD45⁺ DBSCAN clusters across patients. **(L)** Proportion of CD4⁺ and CD8⁺ T cells within CD45⁺ DBSCAN clusters across patients. **(M)** Distribution of RNA_pred_ T cells within CD45⁺ DBSCAN clusters across patients. **(N)** Distributions of *CCR7*-expressing (left) and *GZMB*-expressing (right) T cells within patient 17. **(O)** Proportion of cells expressing selected T cell genes found within versus outside DBSCAN-defined CD45⁺ clusters. **(P)** Spatial distribution of local T cell density in patient 17. **(Q-R)** All genes **(Q)** and T cell-restricted genes **(R)** enriched in high- and low-T cell density regions. Panels show average Spearman correlation coefficients (± error) between gene expression and local T cell density across all patients.

To identify clusters of CD45^+^ immune cells in an unbiased manner, we used Density-Based Spatial Clustering of Applications with Noise (DBSCAN), an unsupervised machine learning algorithm that identifies clusters based on local spatial density^67^. DBSCAN successfully identified regions of high immune cell density, as exemplified in patient 17 (**Figure 6D-E**). In this sample, most DBSCAN clusters were composed predominantly of T cells and myeloid cells, with T cells preferentially localized within clusters and myeloid cells more often dispersed outside of them, consistent with Moran’s I results (**Figure 6F**). Within patient 17’s CD45⁺ aggregates, CD4⁺ T cells outnumbered CD8⁺ T cells, and RNA_pred_-defined T_em_, T_reg_, and Ex1 T cells were enriched among clustered cells (**Figure 6G-H**). The number and size of CD45^+^ DBSCAN clusters varied per patient independent of total immune infiltration (**Figure 6I**, S6E), reflecting spatial heterogeneity in immune organization. Nonetheless, conserved clustering patterns emerged across the cohort, with CD45^+^ niches consistently enriched for T cells and myeloid cells, including higher abundance of CD4^+^ than CD8^+^ T cells, and preferential localization of Ex1, T_reg_, and T_em_ subsets (**Figure 6J-M**).

To leverage our DBSCAN analysis, we examined the spatial distribution of cells expressing key markers of distinct T cell states. Cells expressing naïve and stem-like (*CCR7*, *IL7R*, *SELL, TCF7*), memory-associated (*GZMK*), and regulatory (*FOXP3*) markers were predominantly located within CD45⁺ DBSCAN clusters, while cells expressing activation and exhaustion markers (*HAVCR2*, *ENTPD1*, *GZMA*, *GZMB*) were more frequently found outside of clusters (**Figure 6N-O**), consistent with previous studies^24, 68, 69^. As an orthogonal approach, we analyzed the spatial density of T cells (**Figure 6P**). Genes associated with naïve and lymphoid lineages (*CCR7*, *CD79A*) were enriched in high-density regions, while tumor-associated genes (*MALL*, *SOX2*) were elevated in low-density areas (**Figure 6Q**), suggesting that high T cell density zones reflect lymphoid-like niches and low-density regions are dominated by tumor cells. Within T cells, naïve/memory-like genes (*CCR7*, *SELL*) were enriched in high-density areas, whereas exhaustion- and residence-associated genes (*HAVCR2*, *CD101, ITGAE*) were primarily expressed in low-density regions (**Figure 6R**). These findings further support that exhausted T cells are dispersed across tumor-dense areas, whereas non-exhausted subsets concentrate in immune-dense zones.

### Spatial mapping resolves individual tumor-infiltrating T cell clones at single-cell spatial resolution

While spatial gene expression analyses revealed clear links between T cell phenotype and tissue distribution, they are limited by an inability to distinguish individual T cell clones. Our scRNA-seq data identified PBMC-enriched clonotypes with distinct transcriptional profiles in tumors, suggesting that spatial variation could reflect clonal differences rather than phenotypic or spatial plasticity. This distinction is critical to determine whether exhausted T cells near tumors arise from local differentiation of stem-like cells or represent independent clones with distinct fates.

To directly assess clonal identity *in situ*, we developed a strategy to spatially map individual T cell clones at single-cell resolution by designing custom, patient-specific TCR-targeting probes targeting the variable region of abundant tumor-infiltrating TCRs (**Figure 7A**). Each Xenium assay included TCR probes for at least three different patients, allowing mismatched sample-probe combinations to serve as internal negative controls (**Figure 7B**). Most probes were specifically detected in T cells from the corresponding patient tissue but not in mismatched samples (**Figure 7C-E, S8A-B**), confirming both patient and cell-type specificity.

**Figure 7.**
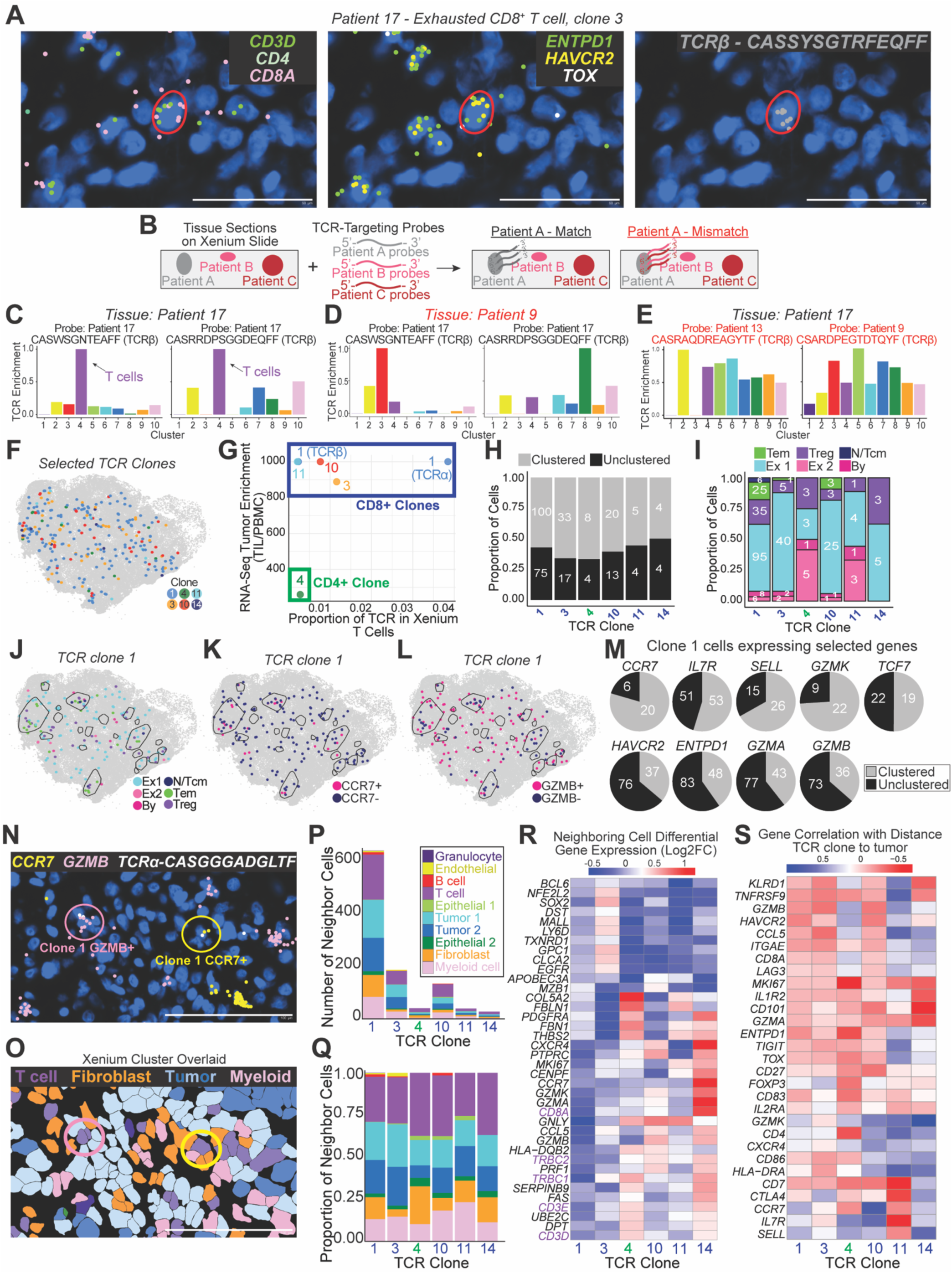
Single-cell spatial TCR mapping detects individual TCR clones *in situ* and identifies phenotype-driven T cell distribution. **(A)** A representative exhausted CD8⁺ T cell visualized in tissue from patient 17, co-expressing *CD3D*, *CD8A*, *ENTPD1*, *HAVCR2*, *TOX*, and a unique TCRβ transcript. Scale bar represents 50 µm. **(B)** Schematic of spatial TCR detection validation workflow using patient-matched and mismatched TCR-targeting probes. **(C-E)** Representative quantification of relative TCR probe enrichment across Xenium gene expression clusters using normalized TCR frequencies for different tissue and probe combinations: patient 17 probe in patient 17 tissue (**C**), patient 17 probe in patient 9 tissue (**D**), and patient 13 probe in patient 17 tissue (**E**). **(F)** Spatial distribution of selected T cell clones in patient 17. Selected clones were most enriched in the Xenium T cell cluster and were detected in a minimum of five T cells. All other cells are shown in gray. **(G)** Tumor enrichment of shown patient 17 TCR clones plotted against its corresponding frequency in the T cell Xenium gene expression cluster. **(H)** Proportion of each selected TCR clone in patient 17 found within and outside of CD45^+^ DBSCAN clusters. **(I)** Proportion of RNA_pred_ cells per selected TCR clone in patient 17. **(J-L)** Spatial location of patient 17 clone 1 cells colored by RNA_pred_ phenotype (**J**), *CCR7* expression (**K**), and *GZMB* expression (**L**) with CD45^+^ DBSCAN clusters outlined over the tissue. **(M)** Number of patient 17 TCR clone 1 cells expressing naïve- and memory-associated (*CCR7, IL7R, SELL*, *GZMK, TCF7*), or exhausted (*HAVCR2, ENTPD1, GZMA, GZMB*) markers found within CD45^+^ DBSCAN-defined immune clusters. **(N-O)** Representative DAPI-stained (**N**) and Xenium cluster-overlaid (**O**) images of patient 17 tissue showing two transcriptionally distinct clone 1 T cells with different neighboring cell types. **(P-Q)** Number (**P**) and proportion (**Q**) of neighbors for each selected clone in patient 17. Neighbors were defined by the five nearest cells within a 15 µm radius. **(R)** Differential gene expression in neighboring cells across selected TCR clones in patient 17. **(S)** Pearson correlation between gene expression within each TCR clone and its distance to the nearest tumor cell.

TCR probes detected in at least five T cells and enriched within the T cell Xenium gene expression cluster were considered successful, yielding 61 validated probes present in over 2,000 T cells in eight patients (**Figure S8C**). We observed a modest correlation between TCR frequencies in scRNA-seq and Xenium (**Figure S8D**). As all probes were pre-selected to fall within a defined range of GC content and melting temperature, detection efficiency showed no association with either parameter (**Figure S8E-F**). To evaluate reproducibility, we profiled seven patient samples in duplicate across two independent Xenium runs. Six of seven samples yielded successful TCR probes in both runs, with substantial overlap in the detected probe sets (**Figure S8G-I**), and variability thus likely reflects differences in tissue content and clonal representation between sectioned regions. Together, these results demonstrate that our patient-specific TCR-targeting probe strategy enables specific and reproducible detection of individual T cell clones at single-cell resolution in human tumors.

### Spatial mapping of T cell clones reveals microenvironment-driven functional states

We next mapped individual T cell clones across the TME to investigate how clonal identity relates to spatial organization and functional state. In patient 17, six tumor-enriched TCR clones were detected, each broadly distributed across the tissue (**Figure 7F-G**). All clones showed a modest preference for localization within CD45⁺ DBSCAN clusters and were largely composed of exhausted RNA_pred_ phenotypes (**Figure 7H-I**).

Focusing on the most abundant CD8^+^ clone (clone 1), we observed clear spatial segregation of RNA_pred_-defined subsets and cells expressing stem-like or exhausted genes. Cells with more plastic states were enriched within CD45⁺ immune clusters, while exhausted cells were more dispersed (**Figure 7J-M**). In one region, two nearby cells from clone 1-one expressing *GZMB* and the other *CCR7* - were surrounded by markedly different neighbors and differed in CD45^+^ DBSCAN spatial clustering. The *GZMB*⁺ cell was surrounded by tumor cells and found outside of a DBSCAN cluster, whereas the *CCR7*⁺ cell was adjacent to myeloid cells and fibroblasts and was located within a CD45^+^ DBSCAN cluster (**Figure 7N-O**). This illustrates not only the spatial resolution of our approach, but also how cells sharing a TCR can adopt divergent phenotypes and occupy distinct tissue microenvironments.

Cross-clonal comparisons within patient 17 revealed subtle differences in spatial context. The only detected CD4^+^ clone (clone 4) was more frequently adjacent to fibroblasts, while CD8^+^ clones varied in neighborhood preferences (**Figure 7P-R**). For example, clone 3 was often near tumor cells (neighbors enriched in *NFE2L2*, *MALL*), whereas clone 14 was more frequently adjacent to other T cells (neighbors enriched in *CD3E*, *CD8A*). Correlations between gene expression and tumor proximity also varied by clone, with clones 1, 3, and 10 showing increased expression of exhaustion markers near tumor cells, and the CD4^+^ clone exhibiting increased *FOXP3* expression in proximity to tumor cells (**Figure 7S**). These findings reveal functional and spatial diversity among individual clones and demonstrate the power of our approach to profile individual T cells *in situ*.

### T cell clones significantly differ in clustering, phenotype, and tumor enrichment across patients

Given the link between organized immune aggregates and T cell differentiation/responses^18, 63, 68, 70–72^, we compared two patients with similar clinical features (p16^+^, smokers, primary tumors) and comparable immune infiltration as measured by flow cytometry (exhausted^high^ CD45^+^ flow group), but with contrasting spatial immune organization - one with a large immune niche (patient 12) and the other with immune cells more dispersed throughout the TME (patient 9). In patient 12, we mapped nine TCR clones, with several identified by paired TCR α and β probes (**Figure 8A-B**). All clones were CD8^+^, primarily found within CD45⁺ DBSCAN clusters, characterized by Ex1 or more plastic T_em_ (potentially representing stem-like cells) RNA_pred_ phenotypes, and were all enriched in the tumor compared to the blood (**Figure 8C-G**). In contrast, we detected ten clones, including both CD4^+^ and CD8^+^ clones, in patient 9 (**Figure 8H-J**). Unlike patient 12, clones in patient 9 were largely found outside of CD45^+^ DBSCAN clusters and were predominantly of the Ex1 RNA_pred_ phenotype, with minimal T_em_-like cells (**Figure 8K-M**). The increased number of stem-like cells in patient 12 may reflect the supportive microenvironment provided by the large immune aggregate within the tumor.

**Figure 8.**
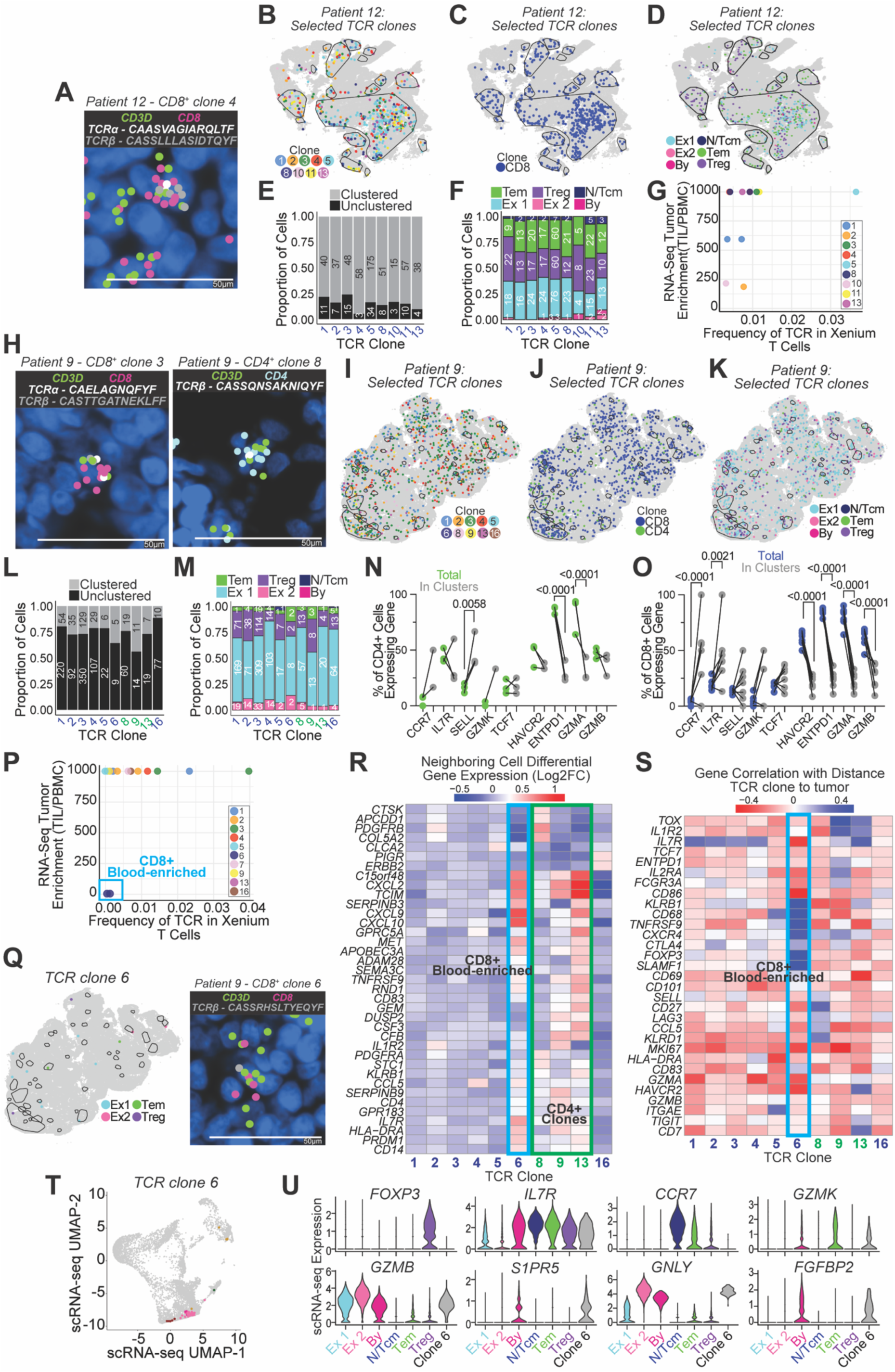
Clonal T cells differ in phenotype and spatial behavior across patients. **(A)** Representative image of a CD8⁺ T cell from patient 12 with detected TCRα- and β-targeting probes. **(B-D)** Spatial distribution of T cell clones within patient 12 tumor tissue (**B**), clone subtypes (**C**), and clonal RNA_pred_ phenotypes (**D**). CD45^+^ DBSCAN clusters are outlined over the tissue. **(E-F)** Proportion of patient 12 TCR clones located within CD45^+^ DBSCAN clusters (**E**) and their assigned RNA_pred_ phenotypes (**F**). **(G)** Tumor enrichment of each detected patient 12 TCR clone plotted against Xenium-detected frequency. **(H)** Representative image showing spatial detection of CD8⁺ (left) and CD4⁺ (right) TCR clones in patient 9. **(I-K)** Spatial distribution of T cell clones within patient 9 tumor tissue (**I**), clone subtypes **(J)**, and clone RNA_pred_ phenotypes **(K)**, with CD45^+^ DBSCAN clusters overlaid. **(L-M)** Proportion of patient 9 TCR clones found within CD45^+^ DBSCAN clusters **(L**) and their assigned RNA_pred_ phenotypes (**M**). **(N-O)** Percentage of CD4⁺ **(N)** and CD8⁺ **(O)** TCR clonal T cells expressing selected genes, shown for all clonal cells (color) and those within CD45^+^ DBSCAN clusters (grey). Each dot pair represents one clone. P values were calculated by mixed-effects analysis with Sidak’s multiple comparisons test. **(P)** Tumor enrichment of each patient 9 clone plotted against its frequency detected by Xenium. **(Q)** Distribution of TCR clone 6, patient 9 throughout the TME (left) and within a representative cell (right). **(R)** Differential gene expression in neighboring cells of patient 9 clones. **(S)** Pearson correlation between gene expression and distance to the nearest tumor cell within T cell clones. **(T)** Location of patient 9, clone 6 TCR in scRNA-seq data, colored by scRNA-seq cluster. **(U)** scRNA-seq gene expression of patient 9, clone 6 cells compared to selected scRNA-seq clusters.

Although stem-like RNA_pred_-defined T cells were less abundant in patient 9, those expressing stem-like markers (*CCR7*, *IL7R*, *SELL*, *TCF7*) were still preferentially located within CD45⁺ DBSCAN clusters, while those expressing exhaustion or cytotoxic genes (*HAVCR2*, *ENTPD1*, *GZMA*, *GZMB*) were enriched outside clusters (**Figure 8N-O**). This trend was consistent across both CD4^+^ and CD8^+^ clones, though more pronounced among CD8⁺ cells. Of note, all clones were highly tumor enriched except for clone 6, which was more abundant in PBMCs than TILs (**Figure 8P**) and may thus be a bystander clone. Despite its low intratumoral frequency, clone 6 was broadly distributed throughout the tumor, showed a distinct neighbor expression profile compared to tumor-enriched clones, and exhibited unique gene-distance correlation patterns (**Figure 8Q-S**). Notably, the blood-enriched clone 6 cells co-localized with influenza A-specific T cells on the scRNA-seq UMAP and transcriptionally resembled the scRNA-seq-defined bystander cluster (**Figure 8T-U**), supporting its potential identity as a bystander clone that has migrated into the tumor.

## Discussion

In this study, we introduce a novel TCR spatial profiling framework that enables *in situ*, single-cell resolution analysis of T cell clonality, transcriptional state, and spatial localization within intact human HNSCC tumors. This framework overcomes longstanding challenges by simultaneously resolving T cell clonality, phenotype, and tissue context. Whereas prior approaches required tissue dissociation or failed to capture clonotype identity, our approach enables *in situ* clonal tracking at single-cell resolution across entire tumor sections.

Stem- and memory-like T cells preferentially localized within immune niches, while exhausted cells were more spatially dispersed throughout tumor-dense regions. These patterns support a model in which proximity to immune-rich hubs preserves more plastic T cell states by providing signals for quiescence and self-renewal, whereas T cells that leave these supportive hubs and migrate towards tumor-dense regions progressively acquire an exhausted phenotype^18, 24, 68, 69^. However, without clonal resolution, such organization could reflect the segregation of unrelated T cell clones - such as bystanders versus tumor-enriched cells - rather than true phenotypic and spatial plasticity. To address this, we mapped individual T cell clones *in situ* by employing patient-specific TCR-targeting probes in spatial transcriptomics. Clonal identity alone did not determine spatial localization. Instead, cells from the same clone often exhibited distinct spatial distributions and transcriptional states, with stem-like or memory cells clustering in immune niches and exhausted members broadly dispersed. These observations indicate that local environmental signals - such as TCR engagement by tumor cells, inhibitory PVR-TIGIT interactions between tumor and T cells, and CD86-mediated signaling between myeloid and T cells - drive the spatial behavior and differentiation of T cells within the TME.

Although local cues may shape most spatial behavior, antigen specificity can also influence localization patterns. A blood-enriched CD8⁺ clone adopted spatial characteristics more typical of CD4^+^ clones rather than tumor-enriched CD8⁺ clones, reinforcing the link between spatial positioning and functional state. Although likely non-tumor-specific, T cells from this clone upregulated some exhaustion-associated genes when in proximity to tumor cells. This supports prior findings that bystander T cells can express inhibitory receptors in tumors^36, 37, 73–75^ and suggests that even non-tumor-specific T cells may acquire partial dysfunction or altered spatial behavior in response to local suppressive signals in the TME. Together, our findings provide *in situ* evidence that local cues within the TME govern T cell differentiation outcomes, rather than cell-intrinsic programming or clonal identity alone.

The extent to which this spatial context predicts therapeutic response remains unclear. Although PD-1 pathway blockade improves survival in some HNSCC patients, the determinants of clinical response remain poorly defined^8–10^. Although patients with an exhausted^high^ flow profile might be expected to respond favorably, CD8^+^ T cell infiltration alone does not reliably predict clinical outcome^17^. Our spatial analyses revealed striking variation in T cell organization across patients that were not explained by immunophenotyping or clinical features. For example, although patients 9, 12, and 13 all had p16^+^ disease and belonged to the exhausted^high^ CD45^+^ flow group, their tumors displayed markedly distinct T cell architectures: patient 9 showed even dispersion of T cells, patient 12 had dense clustering of T cells within a large immune aggregate, and patient 13 exhibited immune exclusion. Thus, even high-parameter flow cytometry, scRNA-seq, and clinical information are not enough to predict T cell behavior within the TME. Instead, differences in spatial architecture may underlie differential treatment responses, as supported by prior studies linking spatial organization with therapeutic efficacy^10, 18–35, 68^.

Interestingly, this spatial architecture correlated with T cell phenotype and plasticity. Patient 12, who exhibited more organized immune structures *in situ*, by scRNA-seq also had lower frequencies of exhausted T cell clusters and increased TCR sharing between T_em_-like and exhausted CD8^+^ T cells, supporting the hypothesis that these regions permit phenotypic plasticity of tumor-specific clones. Consistent with this, Xenium showed that T cells with identical TCRs in patient 12 were more frequently found in plastic states, compared to patients 9 or 13, reinforcing the link between spatial context and phenotypic flexibility. Since immune-rich niches have been associated with improved prognosis and ICB responsiveness^70, 71^, their abundance may further contribute to variability in clinical outcomes.

Together, our multimodal approach establishes a versatile platform for spatially-resolved, antigen-specific T cell profiling in human tumors. By directly detecting endogenous paired TCRs and mapping them alongside gene expression and spatial context, we show that phenotype and specificity influence the spatial distribution of T cells within the HNSCC TME. This plasticity within individual clones underscores the dynamic nature of T cell fate in tumors, shaped continuously by local environmental cues. Our framework provides critical insight into the spatial regulation of T cell exhaustion and therapeutic responsiveness in HNSCC. More broadly, it offers a powerful blueprint for dissecting immune architecture in other solid tumors, with applications in identifying tumor-reactive TCRs, mapping resistance mechanisms, and informing the design of spatially targeted immunotherapies.

### Limitations of the Study

While our findings highlight the importance of spatial organization in shaping T cell function and response to ICB, the resolution and interpretability of our spatial data were limited by the current constraints of spatial transcriptomics. Unlike scRNA-seq, which provides transcriptome-wide resolution of gene expression, Xenium-based spatial transcriptomics retains tissue architecture but in this study was restricted to measuring expression of a 477-gene panel. This limited our ability to fully capture transcriptional diversity and likely contributed to the ∼75% accuracy in predicting scRNA-seq cluster identities using a random forest classifier. Similarly, receptor-ligand interaction analyses such as CellChat were also constrained by gene panel coverage, limiting the detection of potential signaling interactions. Our spatial approach also focused on highly abundant TCRs, potentially missing rare but functionally important clones.

From a biological standpoint, cryopreserved TILs and PBMCs were used for flow cytometry and scRNA-seq, which may underrepresent short-lived or fragile cell populations, although these would be captured in fresh-frozen tissue used for spatial transcriptomics. Additionally, while we observed substantial inter-patient heterogeneity in T cell spatial organization and phenotype, our limited sample size may constrain the full characterization of this variability, and additional spatial patterns may remain undetected. Finally, this study focuses solely on HNSCC, which may limit generalizability to other tumor types with different immune microenvironments or spatial architectures.

## Acknowledgements

We would like to thank Ashley Benham-Duret, Vicky Yao, Patricia Castro, Jeanine van Nostrand, and Joel M. Sederstrom. This work was supported by a Technology Impact Award from the Cancer Research Institute, a Lung Cancer Discovery Award from the American Lung Association, support from the National Cancer Institute (NCI; R37CA285289) and the Adrienne Helis Malvin Medical Research Foundation, and a first-time tenure-track award (RR220069) from the Cancer Prevention and Research Institute of Texas (CPRIT) to W.H.H. W.H.H. is a CPRIT Scholar in Cancer Research. V.C.S. is supported by the NCI through U54CA274321 and by the Veterans Affairs Basic Clinical Science Research and Development division through 1I01CX002776. Tissue acquisition was facilitated by the Human Tissue Acquisition & Pathology Core Baylor College of Medicine. This project was also supported by the Single Cell Genomics Core and Cytometry and Cell Sorting Core at Baylor College of Medicine, with funding from the CPRIT Core Facility Support Award (RP180672) and the NIH (P30CA125123 and S10RR024574). This manuscript does not represent the position, policy, views, or opinions of the National Institutes of Health or the Department of Veterans Affairs.

## Data Availability

PBMC TCR-sequencing data is available from the Gene Expression Omnibus under accession GSE285951. Single-cell RNA-sequencing is available from the Gene Expression Omnibus under accession GSE287301. Xenium data is available from the Gene Expression Omnibus under accession GSE300147.

## Author Contributions

D.J.H. and V.C.S. recruited patients and procured tissue and blood specimens. E.K., S.H., and K.A.M. isolated and cryopreserved PBMCs and TILs. E.K. performed tissue sectioning and K.A.M. performed Xenium spatial transcriptomics assays and analysis and flow cytometry analysis. K.A.M, E.K., S.H., A.Y.X., and C.J.H. performed flow sorting and scRNA-seq library generation. S.H. generated PBMC TCR-sequencing libraries. J.S.L. performed review of Xenium images. A.W. contributed to experimental design and manuscript preparation. W.H.H. analyzed scRNA-seq and PBMC TCR-seq data. K.A.M. and W.H.H. drafted the manuscript with input and review from all authors.

## Methods

### Human samples

Following Institutional Review Board approval (protocol H-40168), HNSCC tumor specimens were obtained from patients treated at two Baylor College of Medicine-affiliated hospitals, Harris Health Ben Taub Hospital and the Michael E. DeBakey Veterans Affairs Medical Center. Tumor specimens were placed in Leibovitz’s L-15 media (Cytiva) and kept at 4 °C until processing, typically within 30 minutes of initial harvest (biopsy or surgical extirpation). Blood samples were collected concurrently in BD Vacutainer EDTA tubes. Healthy donor PBMCs were isolated from buffy coat samples obtained from the Gulf Coast Regional Blood Center. Samples from 28 patients were analyzed in this study; one patient’s pathology was changed from a preliminary diagnosis of HNSCC to ameloblastoma after scRNA-sequencing analysis had been completed. Cells from this patient were used for UMAP projection and clustering but were excluded in all subsequent analyses. Patient, tumor, and treatment characteristics are summarized in **Table S1** and **Table S2**.

### Tissue processing

Tumor samples were collected and processed under aseptic conditions in a biosafety cabinet. Immediately following tumor resection, one portion of the tumor was cryopreserved for downstream spatial transcriptomics by first drying the tumor, embedding it in optimal cutting temperature (OCT) compound in a cryomold, and flash freezing it in a dry ice/isopentane bath before storage in the vapor phase of liquid nitrogen. The remaining tumor tissue was weighed, minced, and digested in 25 mL of Leibovitz’s L-15 medium (Cytiva) supplemented with 250 µL of a 100× tumor enzyme digestion cocktail (a 0.22 µm sterile-filtered solution of 500 mg Collagenase I, 500 mg Collagenase IV, 200 mg DNase I, and 200 mg Elastase in 40 mL of PBS). The sample was shaken for 1 hour at 37 °C and 1000 rpm on an Eppendorf ThermoStat C, then filtered through a 70 µm cell strainer using a 3 mL syringe plunger to ensure thorough dissociation. Cells were washed with additional Leibovitz’s L-15 medium, centrifuged at 500 *g* for 8 minutes, and resuspended in 8 mL of 44% isotonic Percoll diluted in RPMI. A 5 mL volume of 67% isotonic Percoll diluted in PBS was carefully underlaid to form a density gradient, followed by centrifugation without braking at 850 *g* for 20 minutes at room temperature. Tumor-infiltrating lymphocytes (TILs) were collected from the interface, washed with PBS containing 2% FBS and 2 mM EDTA (FACS buffer), and centrifuged at 500 *g* for 8 minutes at 4 °C.

To collect peripheral blood mononuclear cells (PBMCs), 5 mL of whole blood was transferred into a 15 mL conical tube. Using a Pasteur pipette, 5 mL of lymphocyte separation medium (Corning) was added to the bottom of the blood sample to form a gradient. The gradient-containing tube was centrifuged at 850 *g* at room temperature for 20 minutes without braking. The PBMCs at the interface were collected, washed with FACS buffer, and centrifuged at 500 *g* for 8 minutes at 4 °C.

Immediately following TIL and PBMC isolation, the number of CD45^+^ immune cells isolated was counted by flow cytometry with a Cytek Aurora. Samples were then viably cryopreserved by resuspending samples at 1-5 million cells/mL in FBS containing 10% DMSO, placed in a CoolCell freezing chamber at -80 °C overnight, and subsequently transferred to vapor-phase liquid nitrogen storage.

### Cell sorting

Cryopreserved TILs and PBMCs were quickly thawed, transferred to a 15 mL falcon tube, washed with 10 mL of pre-warmed RPMI with 10% FBS, then spun at 500 *g* for 10 min. The resulting pellet was resuspended in FACS buffer, transferred into a 96-well U-bottom plate, and centrifuged at 500 *g* for 10 min at 4 °C. The cell pellets were resuspended in FACS buffer containing Ghost viability dye, anti-human CD16, and biotinylated anti-human TCRv α7.2 (**Table S3**) and incubated on ice for 30 min. Following the first stain, the plate was spun at 500 *g* for 5 min at 4 °C, then the cells were Fc blocked by resuspending in Human TruStain FcX (BioLegend) at 4 °C for 10 min. After blocking, the cells were spun at 500 *g* for 5 min at 4 °C, divided into 4 different hashing groups, then stained with extracellular antibodies and specific hashing antibodies (**Table S3**) in BD Horizon Brilliant Stain Buffer for 30 min on ice. Following incubation, the cells were washed then resuspended in 400 µL of FACS buffer. Samples were sorted using a 5-laser Cytek Aurora Cell Sorter available in the Cytometry and Cell Sorting Core at Baylor College of Medicine.

### Flow cytometry data analysis

Flow cytometry data were analyzed in FlowJo, using the Downsample, FlowSOM, UMAP, and Cluster Explorer plugins. Summary graphs and statistics were generated in GraphPad Prism v10 and RStudio.

### Single-cell RNA-sequencing

Sorted CD45^+^CD3^+^ cells were pooled, mixed with reverse transcription master mix, and loaded into a GEM-X chip according to the manufacturer’s protocol (10x Genomics). Reverse transcription, cDNA amplification, and generation of the gene expression and CITE-seq libraries were also performed according to the manufacturer’s protocol. For TCR sequencing, the manufacturer’s and previously-reported^76^ primer sequences were used to enrich TCRα (outer primer: TGCATGTGCAAACGCCTTCA, inner primer: CGTGTACCAGCTGAGAGACT), TCRβ (outer primer: TCAGGCAGTATCTGGAGTCATTGAG, inner primer: TCTGATGGCTCAAACACAGC), TCRγ (outer primer: TGTGTCGTTAGTCTTCATGGTGTTCC, inner primer: GATCCCAGAATCGTGTTGCTC), and TCRδ (outer primer: AGCTTGACAGCATTGTACTTCC, inner primer: TCCTTCACCAGACAAGCGAC) chains in sequential PCR reactions, using the same forward primer (GATCTACACTCTTTCCCTACACGACGC) in all reactions. Fragmentation and subsequent library preparation steps were completed according to the manufacturer’s protocol. Libraries were sequenced on an Illumina NovaSeq X Plus instrument by Admera Health (South Plainfield, NJ).

### Single-cell RNA-sequencing data analysis: gene expression

Sequenced gene expression, CITE-sequencing, and TCR libraries were analyzed with 10x Genomics Cell Ranger Multi v8.0.1, accessed via 10x Genomics Cloud Analysis. Gene expression and CITE-sequencing reads were assigned to cells with cellranger count, with gene expression reads aligned to the human (GRCh38) genome and assigned to transcripts using the 2024-A reference with Ensembl 110 annotations^77^. Reads mapping to intronic regions were included in gene expression analysis.

Gene expression analysis was performed in Seurat v. 5.1.0^78^. Cells with >80% of CITE-seq reads originating from a single hashing antibody were assigned to a patient sample and moved forward for analysis. Cells with >10% mitochondrial gene content, <200 genes detected, and/or <1000 UMIs were discarded, resulting in a total of 251,708 cells passing quality control. Gene expression was normalized with Seurat’s SCTransform command. Nearest-neighbors were calculated with the FindNeighbors command in Seurat, calculating 100 neighbors with 10 principal components. Clusters were calculated with Seurat’s FindNeighbors command, using the Louvain algorithm with multilevel refinement and a resolution of 0.4 (determined after ***clustree*** analysis^79^). Uniform Manifold Approximation and Projection^80^ (UMAP) was performed with 10 principal components, 100 neighbors, and a minimum distance of 0.

Gene set enrichment analysis was performed with the ***fgsea*** package in R^81^ against the Hallmark, BioCarta cell processes, and Gene Ontology Biological Processes gene sets, accessed through MSigDB^82–84^ in December 2024. For GSEA analysis of each cluster, Seurat’s FindMarkers was used to identify differentially expressed genes between the cluster and all other cells. The sign(log_2_ fold change) * -log_10_(p_adj_) was used as the ranking statistic for the fgsea function.

Graphpad Prism and ggplot2^85^ were used for scRNA-seq data visualization.Graphpad Prism and ggplot2^85^ were used for scRNA-seq data visualization. Raw data for volcano plots are given in **Table S4**.

### Single-cell RNA-sequencing data analysis: Single-cell TCR-sequencing analysis

TCRα/β sequencing reads were assembled and assigned to cells using 10x Genomics’ v7.1.0 TCR reference data and cellranger vdj. Since TCR library preparation also included primers designed to amplify TCRγ/δ sequences, we also performed MiXCR analysis with the mixcr analyze 10x-sc-xcr-vdj pipeline (MiXCR version 4.6.0)^86, 87^. Cells were first assigned a TCR, if available, from cell ranger vdj output. Cells with unassigned TCRs were then assigned a TCR based on MiXCR results.

MAIT cells were identified as those with a TCRα comprised of *TRAV1-2* and *TRAJ33*, *TRAJ20*, or *TRAJ12*^88^. For MAIT-specific phenotypic analysis, cells with a MAIT TCR were compared against all non-MAIT cells with successfully sequenced TCRα/β chains. For bystander cell evaluation, paired TCR α/β information was downloaded from the VDJdb browser, accessed in December 2024. Cells from the scRNA-seq dataset with TCR α and β CDR3 regions that exactly matched VDJdb sequences were marked as bystanders with the annotated antigen specificity.

TCR abundance was analyzed using the TCRβ amino acid CDR3 sequence. Clonotypic diversity was measured using the Shannon index as calculated by the diversity function in the ***vegan*** R package^89^. Clonotypic overlap was assessed using the Morisita-Horn similarity index, derived by subtracting the output of the horn_morisita function from 1, as implemented in the ***abdiv*** R package^90^. For all diversity and overlap analyses, groups or clusters with fewer than 50 TCRs were excluded to avoid unreliable estimates from undersampled repertoires.

### Peripheral blood TCR library generation and sequencing

CD45^+^CD3^+^CD4^+^ cells (CD4^+^ T cells) and CD45^+^CD3^+^CD8^+^ cells (CD8^+^ T cells) from peripheral blood were sorted concurrently with other samples, and RNA and DNA were isolated using the Qiagen AllPrep DNA/RNA Micro kit following the manufacturer’s instructions. TCR-sequencing libraries were generated from isolated RNA with the SMART-Seq Human TCR-seq with UMIs kit (Takara Bio) according to the manufacturer’s protocol. Libraries were sequenced on an Illumina NovaSeq X Plus instrument by Admera Health.

PBMCs were not available from one tumor patient (#24). TCR library generation failed for CD8^+^ T cells from two tumor patients (#21 and #25). CD4^+^ and CD8^+^ TCR libraries were also generated from five healthy donor samples.

### Peripheral blood TCR-sequencing data analysis

Sequenced TCR-seq libraries were analyzed with MiXCR v4.6.0^86, 87^, using the mixcr analyze takara-human-rna-tcr-umi-smartseq pipeline preset and the --assemble-clonotypes-by CDR3 option. Diversity and overlap statistics were calculated as described above for single-cell TCR-seq data, using the TCRβ CDR3 amino acid sequences as unique identifiers for TCR sequences and the UMI-corrected uniqueMoleculeCount for clone abundance. Overall, 22,309,901 TCRβ UMIs were detected (19,228,393 from tumor patients), comprised of 2,903,291 unique TCRβ CDR3 amino acid sequences.

For bulk PBMC analyses, clone abundances were further adjusted to reflect flow cytometry-measured proportions of CD4⁺ and CD8⁺ T cells in each sample. Tumor enrichment for each clone was calculated as the ratio of its TCRβ chain frequency in the tumor to its frequency in matched PBMCs. To avoid inflation from extremely low frequency circulating clones, tumor enrichment values were capped at 1000.

### Xenium Spatial Transcriptomics

Xenium slides were prepared following the Xenium In Situ for Fresh Frozen Tissues User Guide CG000579 from 10x Genomics. 10 µm sections from cryopreserved tissue were placed onto the Xenium slide capture area and stored at -80 °C for up to one week. Slides were subsequently fixed with paraformaldehyde and permeabilized using MeOH as per the manufacturer’s protocol (Xenium In Situ for Fresh Frozen Tissues - Fixation & Permeabilization, User Guide CG000581). Following fixation and permeabilization, ssDNA probe panels were hybridized to the tissue sections overnight, ligated, and amplified by rolling circle amplification prior to cell segmentation staining according to the manufacturer’s protocol (Xenium In Situ Gene Expression with Cell Segmentation Staining, User Guide CG000749). Fluorescent markers included in the staining kit target nuclei (DAPI), membrane boundaries (ATP1A1, CD45, E-Cadherin), interior RNA (18S RNA), and interior proteins (αSMA, Vimentin).

The gene expression in situ hybridization probe set consisted of the predesigned “Human Multi-Tissue and Cancer Panel” (10x Genomics, 377 genes) as well as a custom 100-gene panel targeting 30 key T cell and tumor markers and HPV16 transcripts (**Table S5**) and 70 TCR-targeting probes against a minimum of three patients’ TCRα and TCRβ CDR3 regions. CDR3 regions were selected based on abundance of the target TCR as well as thermodynamic parameters of the probe (i.e., melting temperature and GC content). Data were collected from the prepared slides on a Xenium Analyzer at the Baylor College of Medicine Single Cell Genomics Core using v2.0 analysis, with capture regions selected to cover each tumor section. Immediately following the run, hematoxylin and eosin (H&E) staining was performed on the tissue sections using the manufacturer’s protocol (Xenium In Situ Gene Expression - Post Xenium Analyzer H&E Staining, User Guide CG000613). H&E images were acquired with a Keyence BZ-X810 microscope. Xenium data was processed using the ***Seurat*** package in R and visualized with Seurat, ggplot2, and Xenium Explorer (10x Genomics)^91, 92^.

### Xenium data preprocessing

Data generated from the Xenium Analyzer were imported into the ***Seurat*** package in R^91, 93^. Cells with total gene expression count values equal to zero were removed. Each preprocessed sample was saved as an individual Seurat RDS object for downstream integration.

### Xenium cell type assignment

The analysis included a total of 17 tumor sections derived from 10 unique patient tumors, with technical replicates performed on some patients. Following RDS object import, each Seurat object was loaded individually, normalized using NormalizeData(), and variable features were identified using FindVariableFeatures().

To unify spatial transcriptomics data across all tumor samples and correct for technical variation, anchor-based integration was performed using Seurat’s standard integration workflow. To reduce computational burden for clustering and dimensionality reduction, each dataset was randomly downsampled to a maximum of 20,000 cells. Integration features were selected using SelectIntegrationFeatures() with the number of features set to 500. Integration anchors were identified using FindIntegrationAnchors() with parallel computation enabled via the ***future*** R package^94^. The resulting anchors were used to integrate the datasets with IntegrateData(), yielding a harmonized Seurat object containing batch-corrected gene expression profiles across all 17 sections that preserved biological heterogeneity while minimizing batch effects.

The combined Seurat object was normalized using SCTransform() and scaled using ScaleData(). Uniform Manifold Approximation and Projection (UMAP) and nearest-neighbor graph construction were performed using the top 20 principal components. Gene expression clusters were identified using a clustering resolution of 0.25. Cluster-specific marker genes were identified using Seurat’s FindAllMarkers() function.

Following clustering, cluster labels from the downsampled and integrated dataset were transferred back onto each individual Seurat object by matching cell barcodes. The labeled subset of cells (up to 20,000 per tumor section) was used to extract log-normalized gene expression matrices. These reference datasets were then used to train a random forest classifier using the ***caret*** R package^95^ (train(method = “rf”)), with 5-fold cross-validation (trainControl(method = “cv”, number = 5)) and parallel processing enabled via the ***parallel*** and ***doParallel*** R packages^96^. The model was trained on the transposed expression matrix, using integrated cluster identities as categorical response variables. Once trained, the classifier was applied to the complete SCTransform normalized set of cells detected by Xenium in each sample. This supervised approach enabled propagation of integrated gene expression clusters to the full-resolution spatial transcriptomics data, supporting consistent clustering within each Xenium sample and across all samples.

### T cell-cell type distance calculations

The spatial proximity of individual T cells to other cell types within the tumor microenvironment (TME) was quantified by calculating the minimum Euclidean distance to the nearest neighboring cell of a given type. Spatial coordinates for both T cells and comparison cell types were extracted from the Seurat object and converted into spatial point pattern objects using the ***spatstat*** R package^97^. A shared spatial window was defined based on the combined x- and y-coordinate ranges of all cells to ensure alignment. The nncross() function was used to compute the shortest distances, and parallel computation was implemented using the ***future*** R package^94^. Skewness was computed for distance distributions using the skewness() function in R under the ***moments*** package^98^.

### Correlation of gene expression with spatial proximity

Gene-level correlation analysis was then performed to identify transcriptional programs associated with spatial localization. For each gene, a Pearson correlation test was conducted between its expression across T cells and the distance of each cell to the nearest comparison cell type. P-values were adjusted for multiple testing using the Benjamini-Hochberg false discovery rate (FDR) method. A composite statistic was computed by multiplying the signed correlation coefficient by the negative log_10_ of the adjusted p-value, reflecting the direction of correlation and statistical significance.

To assess reproducible spatial expression patterns across patients, per-sample correlation results were aggregated into a unified dataset. Pearson correlations were first transformed into Fisher Z-scores using the fisherz() function from the ***pysch*** R package^99^, summed across patients on a per-gene basis, and then back-transformed to obtain overall correlation estimates. Fisher’s method was used to combine p-values across samples, and the standard deviation of Z-scores was calculated to quantify inter-sample variability. Genes with consistent positive or negative correlations (combined p_adj_ < 0.05) were defined as spatially up- or downregulated with respect to each target cell type. Correlation results across different cell types were merged into a single dataset, row-scaled (z-score normalization per gene), and visualized as a clustered heatmap generated using ggplot2.

### Nearest-neighbor calculations

Nearest-neighbor relationships were computed using Euclidean distance between cell centroids. For each cell in a given subset (Xenium gene expression cluster, RNA-seq cluster, TCR clone), the five nearest neighboring cells within a 15 µm threshold were identified using a k-nearest-neighbor (kNN) search implemented via the nn2() function from the ***RANN*** R package^100^.

### Assessment of neighbor composition

Following nearest-neighbor identification, the proportion of neighboring cells belonging to each Xenium-predicted gene expression cluster was quantified for every cell subset (Xenium gene expression cluster, RNA-seq cluster, TCR clone). These neighborhood compositions were aggregated per patient and visualized as stacked bar plots to compare the local microenvironmental contexts.

### Moran’s I spatial autocorrelation analysis

Spatial neighbors were defined using a k-nearest neighbors algorithm (k = 10) implemented via the knearneigh() and knn2nb() functions from the ***spdep*** R package^66^. Binary vectors indicating group membership (1 = in group, 0 = not in group) were constructed, and Moran’s I was computed using moran.test() from ***spdep***. To evaluate significance, 500 random cell sets were generated for each population by sampling an equal number of cells from the full dataset. Moran’s I was calculated for each random set to create a null distribution for comparison. This analysis was applied across all patient samples for major immune populations, RNA-defined T cells, and transcriptional T cell subsets.

### DBSCAN spatial analysis

Density-based spatial clustering of applications with noise (DBSCAN) was used to identify spatial clusters of distinct cell populations across tissue sections. DBSCAN clustering was performed on the spatial coordinates of cells of the indicated populations using the dbscan() function from the ***dbscan*** package in R^67^, with uniform parameters applied across all samples and cell types: a fixed neighborhood radius (eps) of 80 µm and a minimum of 35 neighboring cells (minPts) to define a cluster. Cells meeting these density criteria were assigned DBSCAN cluster labels, while cells outside of DBSCAN clusters were classified as unclustered.

### CellChat analysis

Cell-cell communication analysis was performed using the ***CellChat*** R package^58^ on the integrated Xenium dataset. The Seurat object was converted to a CellChat object using Xenium gene expression clusters to define cell groups. Signaling interactions were inferred using the CellChat human ligand-receptor database (CellChatDB.human). Overexpressed ligands and receptors were identified with subsetData(), identifyOverExpressedGenes(), and identifyOverExpressedInteractions() based on intra and intercluster gene expression. Communication probabilities were calculated using computeCommunProb(), with interactions involving fewer than five cells excluded. Pathway-level signaling was computed with computeCommunProbPathway() and aggregated using aggregateNet(). Signaling roles were evaluated using netAnalysis_computeCentrality(), and networks were visualized with netVisual_circle(), netVisual_aggregate(), and netVisual_heatmap(). Pathway summaries were used to identify dominant sending and receiving clusters.

### TCR probe analysis

To evaluate TCR probe enrichment across transcriptionally defined T cell clusters, we identified cells with detectable expression of each TCR probe (counts >0) and calculated their frequency within each Xenium gene expression cluster. For each probe, frequencies were normalized to the total number of cells per cluster and then scaled such that the cluster with the highest enrichment was assigned a value of 1. This allowed for relative enrichment comparisons across clusters on a per-probe basis. Probes detected in at least five cells and showing peak enrichment in a T cell cluster were classified as successful. Enrichment values for both matched and mismatched probes were visualized using ggplot2.

### Prediction of scRNA-seq clusters (RNA_pred_) from Xenium data

To transfer T cell subtype annotations from scRNA-seq to spatial transcriptomics data generated with the Xenium platform, we trained a supervised classification model using a random forest algorithm. Genes for use in modeling were selected based on expression in T cells across both platforms, excluding those expressed by tumor cells to minimize spurious signal from segmentation or transcript localization artifacts. This filtering yielded a panel of 52 genes used for label transfer. We also excluded quiescent and proliferating T cell clusters from model training, as these states are harder to assess in spatial data and were often confounded by transcript spillover from highly transcriptionally active dividing tumor cells. The resulting model was trained to classify cells into seven T cell clusters (7, 8, 10, 11, 12, 13, and 14). Expression values from scRNA-seq data were log-transformed followed by min-max scaling within each gene to generate a percent-of-maximum expression matrix. Cells were randomly partitioned into training (80%) and validation (20%) subsets, and a random forest classifier with 500 trees was trained using the ***ranger*** package in R^101^ to predict T cell subtypes. The most frequent cluster (cluster 10) comprised 26.1% of the training dataset, providing a minimum baseline for evaluating model performance; the random forest classifier achieved 75.6% accuracy on held-out validation cells.

To apply the trained classifier to Xenium data, gene expression values of cells within the T cell Xenium gene expression cluster were log-transformed and scaled to percent of maximum, matching the preprocessing steps used for model training. The random forest classifier was then applied to assign each Xenium T cell to its most probable RNA-seq-defined T cell subtype (RNA_pred_). These predicted labels were retained for all downstream analyses and spatial visualizations.

To explore whether correcting for differences in cell subtype proportions across platforms would improve label transfer, we implemented an additional reweighting step using optimal transport. Although the method successfully enforced proportional alignment with the reference clusters, it often introduced noise or overcorrection and did not improve downstream analyses. We also evaluated multinomial logistic regression as a baseline classifier, which performed reasonably but less accurately than random forest classifiers (71-72% accuracy depending on normalization). Random forest classifiers showed higher accuracy on held-out scRNA-seq cells, including 75.5% accuracy using SCTransform-normalized data. We ultimately selected log-transformed percent-of-maximum normalization for downstream predictions, as it provided comparable accuracy and more effectively accounted for gene-specific differences in probe efficiency in Xenium data.

## Notes

### Competing Interest Statement

The authors have declared no competing interest.

