## Supplementary Figures for "Single-cell, Spatially-Resolved TCR Profiling Links T Cell Phenotype and Clonality in Human Tumors"

[1] Department of Molecular and Cellular Biology, Baylor College of Medicine, Houston, TX, USA
[2] Department of Laboratory Medicine and Pathology, Mayo Clinic, Scottsdale, AZ, USA
[3] Department of Otolaryngology, Pelotonia Institute for Immuno-Oncology, The Ohio State University Comprehensive Cancer Center, College of Medicine, The Ohio State University Wexner Medical Center, Columbus, OH, USA
[4] Bobby R. Alford Department of Otolaryngology Head and Neck Surgery, Baylor College of Medicine, Houston, TX, USA
[5] ENT Section, Operative CareLine, Michael E. DeBakey Veterans Affairs Medical Center, Houston, TX, USA
[6] Center for Translational Research on Inflammatory Diseases, Michael E. DeBakey Veterans Affairs Medical Center, Houston, TX, USA
[7] Center for Cell and Gene Therapy, Baylor College of Medicine, Houston, TX, USA
[8] Dan L. Duncan Comprehensive Cancer Center, Baylor College of Medicine, Houston, TX 77030
[9] These authors contributed equally

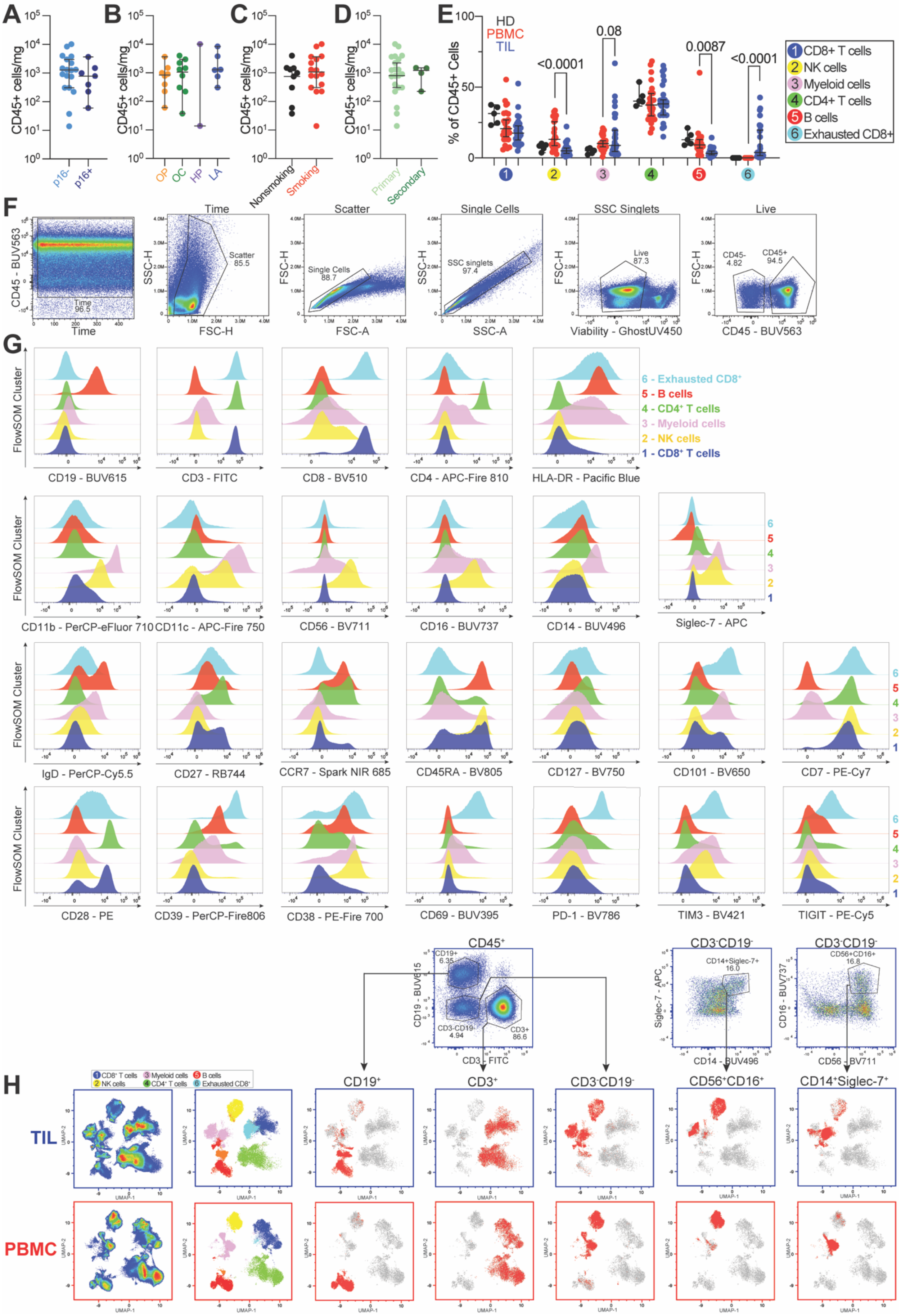

A
B
C
D
E
F
G
H
CD45+ cells/mg
p16-
p16+
OP
OC
HP
LA
Nonsmoking
Smoking
Primary
Secondary
% of CD45+ Cells
HD
PBMC
TIL
<0.0001
0.08
0.0087
<0.0001
1 CD8+ T cells
2 NK cells
3 Myeloid cells
4 CD4+ T cells
5 B cells
6 Exhausted CD8+
Time
Scatter
Single Cells
SSC Singlets
Live
CD45 - BUV563
SSC-H
FSC-H
FSC-A
SSC-A
Viability - GhostUV450
FlowSOM Cluster
CD19 - BUV615
CD3 - FITC
CD8 - BV510
CD4 - APC-Fire 810
HLA-DR - Pacific Blue
CD11b - PerCP-eFluor 710
CD11c - APC-Fire 750
CD56 - BV711
CD16 - BUV737
CD14 - BUV496
Siglec-7 - APC
IgD - PerCP-Cy5.5
CD27 - RB744
CCR7 - Spark NIR 685
CD45RA - BV805
CD127 - BV750
CD101 - BV650
CD7 - PE-Cy7
CD28 - PE
CD39 - PerCP-Fire806
CD38 - PE-Fire 700
CD69 - BUV395
PD-1 - BV786
TIM3 - BV421
TIGIT - PE-Cy5
CD45+
CD3-CD19-
CD19+
CD3+
CD56+CD16+
CD14+Siglec-7+
TIL
PBMC
UMAP-1
UMAP-2

**Figure S1. Flow cytometry staining and conventional gating validates identification of CD45$^+$ FlowSOM clusters.**

(**A-D**) Quantification of CD45$^+$ TILs per milligram of tumor tissue stratified by p16 status (**A**), tumor site (**B**), smoking status (**C**), and tumor recurrence (**D**) reveals no significant differences in immune infiltration across of clinical subgroups. Data represent TILs isolated from 27 HNSCC patients and are shown as median with 95% confidence intervals (CI). Statistical comparisons were performed using the Mann-Whitney test for binary variables (**A, C, D**) and Kruskal-Wallis test with Dunn's multiple comparisons for multi-group comparison (**B**). All P values in panels A-D were non-significant (P>0.05).

(**E**) Distribution of FlowSOM-defined CD45$^+$ populations across healthy donor PBMCs (HD, n = 5), HNSCC patient PBMCs (n = 26), and TILs (n = 27) shows that myeloid cells and exhausted CD8$^+$ T cells are enriched in TILs and NK cells and B cells are more predominant in PBMCs. No differences are observed between HDs and HNSCC patient PBMCs. Twenty-six PBMC-TIL pairs were matched. Error bars are plotted as median with 95% CI. Paired mixed-effects analysis with Sidak's multiple comparisons test was performed on patient-matched PBMC and TIL samples. HD PBMCs vs HNSCC PBMCs were compared using two-way ANOVA with Sidak's multiple comparisons test.

(**F**) Representative gating strategy for identification of live CD45$^+$ immune cells. Time gating was applied to exclude periods of unstable flow. Doublets and aggregates were removed using sequential FSC-A versus FSC-H and SSC-A versus SSC-H plots. Live cells were identified using the viability dye Ghost UV450, and CD45$^+$ events were subsequently gated to isolate the immune cell population.

(**G**) Expression of individual proteins in CD45$^+$ TILs, stratified by FlowSOM-defined immune cluster.

(**H**) Validation of UMAP-based cell type classification using conventional gating. Shown are representative TIL (top) and PBMC (bottom) samples. CD45$^+$ cells were manually gated to identify B cells (CD19$^+$), T cells (CD3$^+$), or non-B/T cells (CD3$^-$CD19$^-$). Within the non-B/T fraction, NK cells (CD56$^+$CD16$^+$) and myeloid cells (CD14$^+$Siglec7$^+$) were identified. Manually gated populations and FlowSOM-defined populations occupy the same regions in UMAP space.

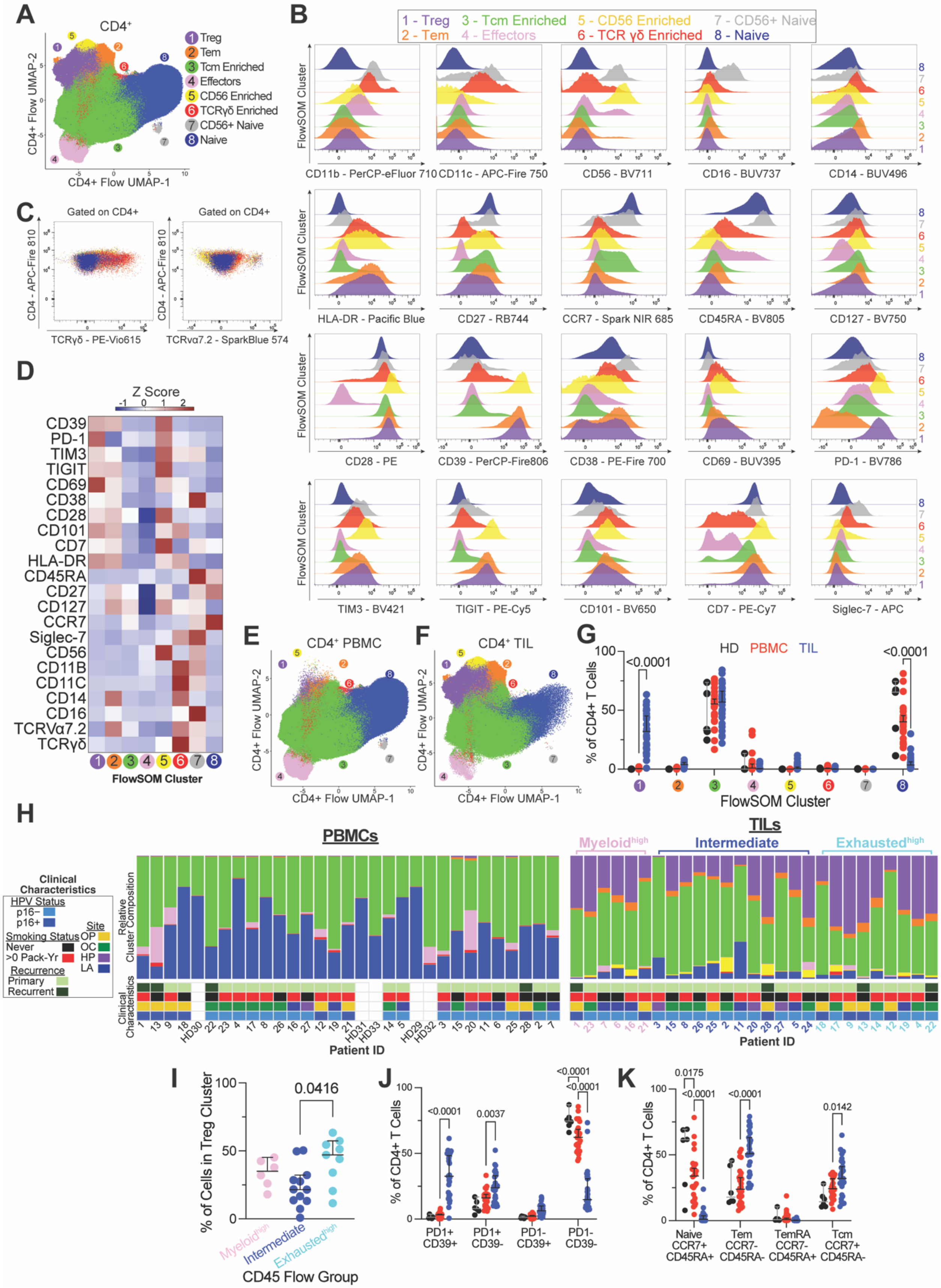

A
CD4+
1 Treg
2 Tem
3 Tcm Enriched
4 Effectors
5 CD56 Enriched
6 TCRγδ Enriched
7 CD56+ Naive
8 Naive
CD4+ Flow UMAP-2
CD4+ Flow UMAP-1
B
1 - Treg
2 - Tem
3 - Tcm Enriched
4 - Effectors
5 - CD56 Enriched
6 - TCR γδ Enriched
7 - CD56+ Naive
8 - Naive
FlowSOM Cluster
CD11b - PerCP-eFluor 710
CD11c - APC-Fire 750
CD56 - BV711
CD16 - BUV737
CD14 - BUV496
HLA-DR - Pacific Blue
CD27 - RB744
CCR7 - Spark NIR 685
CD45RA - BV805
CD127 - BV750
CD28 - PE
CD39 - PerCP-Fire806
CD38 - PE-Fire 700
CD69 - BUV395
PD-1 - BV786
TIM3 - BV421
TIGIT - PE-Cy5
CD101 - BV650
CD7 - PE-Cy7
Siglec-7 - APC
C
Gated on CD4+
CD4 - APC-Fire 810
TCRγδ - PE-Vio615
TCRvα7.2 - SparkBlue 574
D
Z Score
CD39
PD-1
TIM3
TIGIT
CD69
CD38
CD28
CD101
CD7
HLA-DR
CD45RA
CD27
CD127
CCR7
Siglec-7
CD56
CD11B
CD11C
CD14
CD16
TCRVα7.2
TCRγδ
FlowSOM Cluster
E
CD4+ PBMC
F
CD4+ TIL
G
HD PBMC TIL
% of CD4+ T Cells
<0.0001
FlowSOM Cluster
H
PBMCs
TILs
Myeloid^high
Intermediate
Exhausted^high
Clinical Characteristics
HPV Status
p16−
p16+
Site
OP
OC
HP
LA
Smoking Status
Never
>0 Pack-Yr
Recurrence
Primary
Recurrent
Relative Cluster Composition
Patient ID
I
% of Cells in Treg Cluster
0.0416
Myeloid^high
Intermediate
Exhausted^high
CD45 Flow Group
J
% of CD4+ T Cells
<0.0001
0.0037
<0.0001
<0.0001
PD1+ CD39+
PD1+ CD39-
PD1- CD39+
PD1- CD39-
K
% of CD4+ T Cells
0.0175
<0.0001
<0.0001
0.0142
Naive CCR7+ CD45RA+
Tem CCR7- CD45RA-
TemRA CCR7- CD45RA+
Tcm CCR7+ CD45RA-

**Figure S2. Flow cytometry analysis of circulating and tumor-infiltrating $CD4^+$ T cells.**

(**A**) Manually gated $CD4^+$ T cells from healthy donor PBMCs (HD, n = 5), HNSCC patient PBMCs (PBMC, n = 26), and HNSCC TILs (n = 27) were concatenated and clustered using FlowSOM. UMAP shows $CD4^+$ T cells from all samples, colored by FlowSOM-defined cluster identity.

(**B**) Flow cytometry staining of $CD4^+$ TILs for individual markers across each FlowSOM-defined $CD4^+$ T cell cluster.

(**C**) Staining of γδ (left) and MAIT (TCRv $\alpha 7.2^+$, right) $CD4^+$ T cell subsets by flow cytometry. Cells are colored by FlowSOM-defined $CD4^+$ cluster. γδ T cells are predominantly found within the TCRγδ-enriched cluster, while MAIT T cells (TCRv $\alpha 7.2^+$) are infrequent and not restricted to a single phenotypic cluster.

(**D**) Heatmap showing relative protein expression across FlowSOM-defined $CD4^+$ T cell clusters further supports cluster identification.

(**E-F**) UMAP projections of $CD4^+$ T cells separated by compartment reveal differences in cluster distribution in PBMCs (**E**) compared to TILs (**F**).

(**G**) Proportion of $CD4^+$ FlowSOM clusters in HDs (n = 5), HNSCC patient PBMCs (n = 26), and TILs (n = 27) reveals significantly more $CD4^+$ $T_{regs}$ in TILs and more naïve $CD4^+$ T cells in PBMCs and HDs. 26 PBMC-TIL pairs were matched. Error bars are plotted as median with 95% CI. Paired mixed-effects analysis with Sidak's multiple comparisons test was performed on patient-matched PBMC and TIL samples. HD PBMCs were compared to HNSCC patient PBMCs analysed using two-way ANOVA with Sidak's multiple comparisons test.

(**H**) FlowSOM-defined $CD4^+$ T cell cluster composition in PBMCs (left) and TILs (right) per patient. Patients are arranged in $CD45^+$ hierarchical clustering order (**Figure 1F**) to show trends between $CD4^+$ clusters and $CD45^+$ flow groups. Visually, $T_{regs}$ appear more abundant in the exhausted$^{high}$ group, while the myeloid$^{high}$ and intermediate groups show no clear differences in $CD4^+$ cluster composition. Clinical variables are annotated below each patient. Tumor sites include oropharynx (OP), oral cavity (OC), hypopharynx (HP), and larynx (LA). Smoking is measured in pack years (Pack-Yr).

(**I**) Percentage of cells found within the $CD4^+$ $T_{reg}$ cluster stratified by CD45 flow group shows $T_{regs}$ were enriched in the myeloid$^{high}$ and exhausted$^{high}$ $CD45^+$ flow groups, with the latter exhibiting the greatest accumulation. P values were calculated using the Kruskal-Wallis test.

(**J**) Proportion of $CD4^+$ T cells present within conventionally gated PD-1/CD39 subsets across HD PBMCs (n = 5), HNSCC patient PBMCs (n = 26), and HNSCC TILs (n = 27). More exhausted/regulatory subsets ($PD1^+$ $CD39^+$ and $PD1^+$ $CD39^-$) are more abundant in TILs and less exhausted ($PD1^-$ $CD39^-$) cells are more abundant in PBMCs. P-values were calculated using paired mixed-effects analysis with Sidak's multiple comparisons test.

(**K**) Proportion of $CD4^+$ T cells present within conventionally gated CCR7/CD45RA subsets across HD PBMCs (n = 5), HNSCC patient PBMCs (n = 26), and HNSCC TILs (n = 27). Naïve cells ($CCR7^+$ $CD45RA^+$) are enriched in PBMCs, while effector memory ($T_{em}$, $CCR7^-$ $CD45RA^-$) and central memory ($T_{cm}$, $CCR7^+$ $CD45RA^-$) T cells are enriched in TILs. P values were calculated using paired mixed-effects analysis with Sidak's multiple comparisons test.

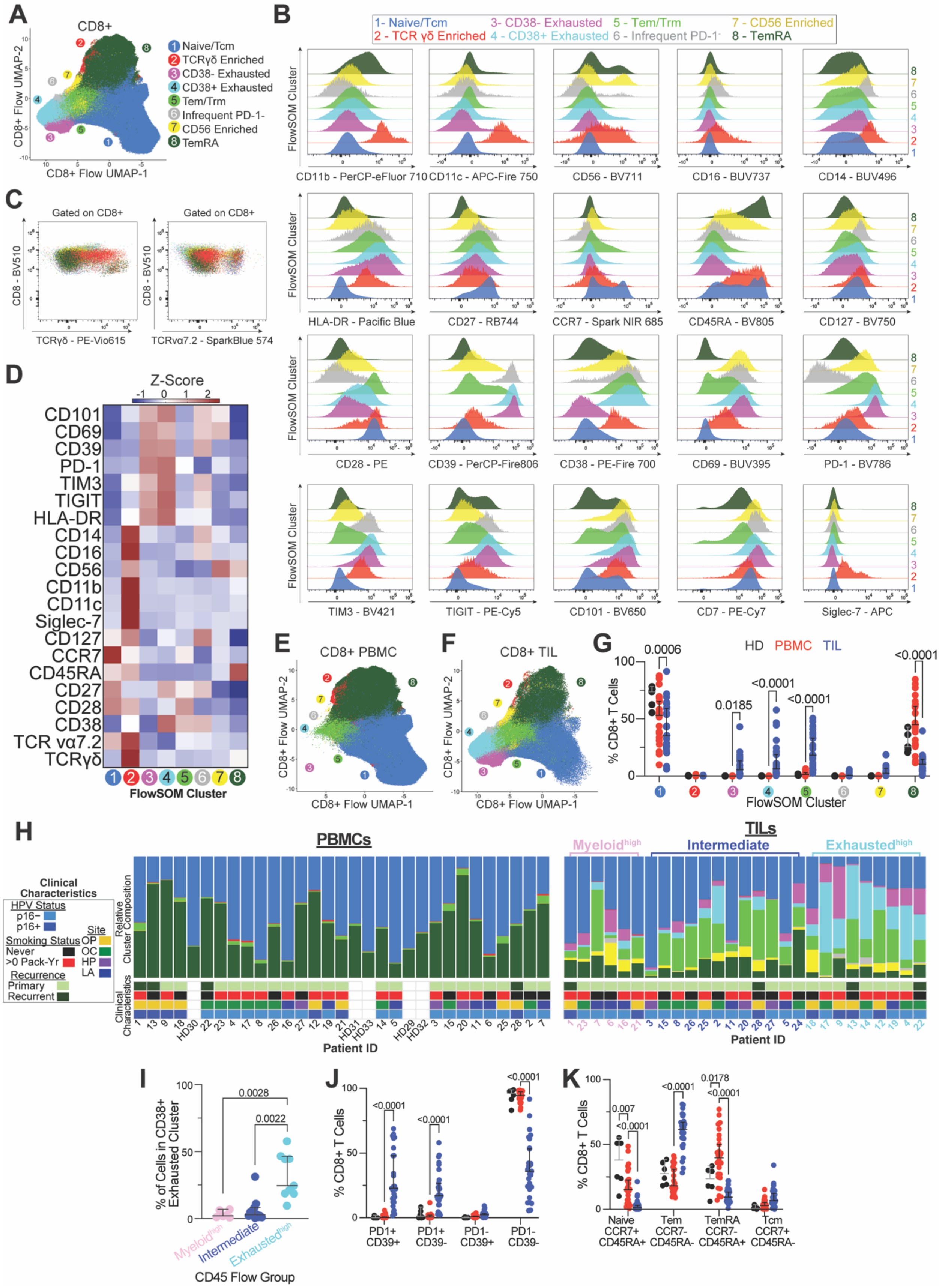
A
CD8+
CD8+ Flow UMAP-2
CD8+ Flow UMAP-1
1 Naive/Tcm
2 TCRγδ Enriched
3 CD38- Exhausted
4 CD38+ Exhausted
5 Tem/Trm
6 Infrequent PD-1-
7 CD56 Enriched
8 TemRA
B
1- Naive/Tcm
2 - TCR γδ Enriched
3- CD38- Exhausted
4 - CD38+ Exhausted
5 - Tem/Trm
6 - Infrequent PD-1-
7 - CD56 Enriched
8 - TemRA
FlowSOM Cluster
CD11b - PerCP-eFluor 710
CD11c - APC-Fire 750
CD56 - BV711
CD16 - BUV737
CD14 - BUV496
HLA-DR - Pacific Blue
CD27 - RB744
CCR7 - Spark NIR 685
CD45RA - BV805
CD127 - BV750
CD28 - PE
CD39 - PerCP-Fire806
CD38 - PE-Fire 700
CD69 - BUV395
PD-1 - BV786
TIM3 - BV421
TIGIT - PE-Cy5
CD101 - BV650
CD7 - PE-Cy7
Siglec-7 - APC
C
Gated on CD8+
CD8 - BV510
TCRγδ - PE-Vio615
TCRvα7.2 - SparkBlue 574
D
Z-Score
CD101
CD69
CD39
PD-1
TIM3
TIGIT
HLA-DR
CD14
CD16
CD56
CD11b
CD11c
Siglec-7
CD127
CCR7
CD45RA
CD27
CD28
CD38
TCR vα7.2
TCRγδ
FlowSOM Cluster
E
CD8+ PBMC
F
CD8+ TIL
G
HD PBMC TIL
% CD8+ T Cells
0.0006
0.0185
<0.0001
<0.0001
<0.0001
FlowSOM Cluster
H
PBMCs
TILs
Myeloid^high
Intermediate
Exhausted^high
Clinical Characteristics
HPV Status
p16−
p16+
Smoking Status
Never
>0 Pack-Yr
Recurrence
Primary
Recurrent
Site
OP
OC
HP
LA
Relative Cluster Composition
Clinical Characteristics
Patient ID
I
% of Cells in CD38+ Exhausted Cluster
0.0028
0.0022
Myeloid^high
Intermediate
Exhausted^high
CD45 Flow Group
J
% CD8+ T Cells
<0.0001
<0.0001
<0.0001
PD1+ CD39+
PD1+ CD39-
PD1- CD39+
PD1- CD39-
K
% CD8+ T Cells
0.007
<0.0001
<0.0001
0.0178
<0.0001
Naive CCR7+ CD45RA+
Tem CCR7- CD45RA-
TemRA CCR7- CD45RA+
Tcm CCR7+ CD45RA-

**Figure S3. Flow cytometry analysis of circulating and tumor-infiltrating CD8$^+$ T cells.**

(**A**) Manually gated CD8$^+$ T cells from healthy donor PBMCs (HD, n = 5), HNSCC patient PBMCs (PBMC, n = 26), and HNSCC TILs (n = 27) were concatenated and clustered using FlowSOM. UMAP shows CD8$^+$ T cell cells from all samples, colored by FlowSOM-defined cluster identity.

(**B**) Flow cytometry staining of individual markers in CD8$^+$ TILs across each FlowSOM-defined CD8$^+$ T cell cluster.

(**C**) Flow cytometry staining for CD8$^+$ γδ and MAIT (TCRv α7.2$^+$) T cell subsets, with cells colored by CD8$^+$ FlowSOM clusters. γδ cells are confined to the TCRγδ enriched FlowSOM cluster, while MAIT cells (TCRv α7.2$^+$) are dispersed across multiple clusters.

(**D**) Heatmap showing relative protein expression across FlowSOM-defined CD8$^+$ T cell clusters reveals differences in protein expression across clusters and supports FlowSOM cluster classification.

(**E-F**) CD8$^+$ T cell UMAP projections from (**A**) split by compartment to reveal differential CD8$^+$ T cell cluster distribution in PBMCs (**E**) and TILs (**F**).

(**G**) Proportion of CD8$^+$ FlowSOM clusters in HDs (n = 5), PBMCs (n = 26), and TILs (n = 27). CD38$^-$ exhausted, CD38$^+$ exhausted, and memory-like clusters are significantly enriched in TILs, whereas naïve and $T_{emRA}$ clusters are more abundant in PBMCs than in TILs. The two exhausted populations (CD38$^{-/+}$) may represent two different states of exhaustion with reduced cytotoxicity within the CD38$^+$ T cells. 26 PBMC-TIL pairs were patient-matched. Error bars are plotted as median with 95% CI. Mixed-effects analysis with Sidak's multiple comparisons test was performed on paired PBMC and TIL samples. HD and HNSCC patient PBMCs were compared using two-way ANOVA with Sidak's multiple comparisons test.

(**H**) FlowSOM-defined CD8$^+$ T cell cluster composition per patient separated by tissue with PBMCs (left) and TILs (right). Patients are ordered by CD45 hierarchical clustering (**Figure 1F**) to show trends across CD45 flow groups. CD8$^+$ exhausted clusters appear enriched in the exhausted$^{high}$ group, naïve/central memory cells appear more abundant in the myeloid$^{high}$ group, and the intermediate group appears to have increased representation of cells within the Tem/Trm (effector and resident memory) cluster. Clinical variables are annotated below each patient. Tumor sites include oropharynx (OP), oral cavity (OC), hypopharynx (HP), and larynx (LA). Smoking is measured in pack years (Pack-Yr).

(**I**) Percentage of cells found within the CD38$^+$ exhausted CD8$^+$ T cell cluster stratified by CD45 flow group shows significantly increased CD38$^+$ exhausted cells in the exhausted$^{high}$ flow group. P values were calculated using a Kruskal-Wallis test.

(**J**) Proportion of CD8$^+$ T cells present within conventionally gated PD-1/CD39 subsets across HD PBMCs (n = 5), HNSCC patient PBMCs (n = 26), and HNSCC TILs (n = 27). More exhausted/regulatory subsets (PD1$^+$ CD39$^+$ and PD1$^+$ CD39$^-$) are more abundant in TILs, while less exhausted (PD1$^-$ CD39$^-$) cells are more abundant in PBMCs. P-values were calculated using paired mixed-effects analysis with Sidak's multiple comparisons test.

(**K**) Proportion of CD8$^+$ T cells present within conventionally gated CCR7/CD45RA subsets across HD PBMCs (n = 5), HNSCC patient PBMCs (n = 26), and HNSCC TILs (n = 27). Naïve cells (CCR7$^+$ CD45RA$^+$) and $T_{emRA}$ cells (CCR7$^-$ CD45RA$^+$) are enriched in PBMCs, while effector memory cells ($T_{em}$, CCR7$^-$ CD45RA$^-$) are enriched in TILs. Unlike CD4$^+$ TILs, minimal CD8$^+$ $T_{cm}$ cells were observed within TILs. P values were calculated using paired mixed-effects analysis with Sidak's multiple comparisons test.

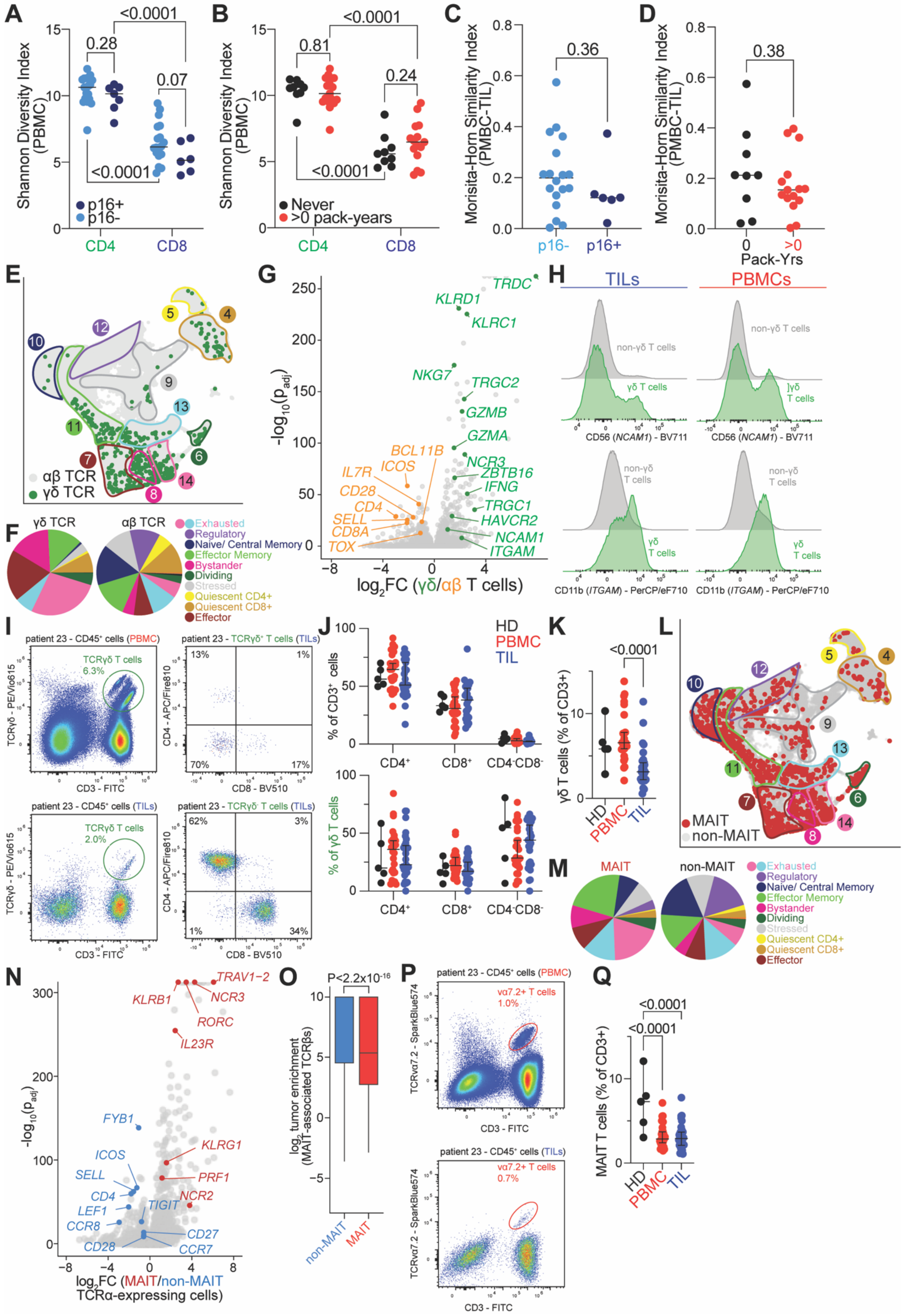

A
B
C
D
Shannon Diversity Index (PBMC)
<0.0001
0.28
0.07
<0.0001
p16+
p16-
CD4
CD8
0.81
<0.0001
0.24
Never
>0 pack-years
Morisita-Horn Similarity Index (PMBC-TIL)
0.36
p16-
p16+
0.38
Pack-Yrs
E
αβ TCR
γδ TCR
F
γδ TCR
αβ TCR
Exhausted
Regulatory
Naive/ Central Memory
Effector Memory
Bystander
Dividing
Stressed
Quiescent CD4+
Quiescent CD8+
Effector
G
TRDC
KLRD1
KLRC1
NKG7
TRGC2
GZMB
GZMA
BCL11B
IL7R
ICOS
NCR3
ZBTB16
CD28
IFNG
CD4
TRGC1
SELL
HAVCR2
CD8A
NCAM1
TOX
ITGAM
log2FC (γδ/αβ T cells)
H
TILs
PBMCs
non-γδ T cells
γδ T cells
CD56 (NCAM1) - BV711
CD11b (ITGAM) - PerCP/eF710
I
patient 23 - CD45+ cells (PBMC)
TCRγδ T cells 6.3%
patient 23 - TCRγδ+ T cells (TILs)
patient 23 - CD45+ cells (TILs)
TCRγδ T cells 2.0%
patient 23 - TCRγδ- T cells (TILs)
TCRγδ - PE/Vio615
CD3 - FITC
CD4 - APC/Fire810
CD8 - BV510
J
HD
PBMC
TIL
% of CD3+ cells
% of γδ T cells
CD4+
CD8+
CD4-CD8-
K
<0.0001
γδ T cells (% of CD3+)
L
MAIT
non-MAIT
M
MAIT
non-MAIT
N
TRAV1-2
KLRB1
NCR3
RORC
IL23R
FYB1
KLRG1
ICOS
SELL
PRF1
CD4
NCR2
LEF1
TIGIT
CCR8
CD27
CD28
CCR7
log2FC (MAIT/non-MAIT TCRα-expressing cells)
O
P<2.2x10^-16
log2 tumor enrichment (MAIT-associated TCRβs)
non-MAIT
MAIT
P
patient 23 - CD45+ cells (PBMC)
vα7.2+ T cells 1.0%
patient 23 - CD45+ cells (TILs)
vα7.2+ T cells 0.7%
TCRvα7.2 - SparkBlue574
CD3 - FITC
Q
<0.0001
<0.0001
MAIT T cells (% of CD3+)

**Figure S4. Characterization of conventional and unconventional T cells in circulation and tumors.**

**(A-B)** Shannon diversity index of circulating $CD4^+$ and $CD8^+$ T cells grouped by p16 (**A**) and smoking status (**B**) reveals lower diversity among $CD8^+$ cells compared to $CD4^+$ cells, regardless of p16 or smoking status.

**(C-D)** Morisita-Horn index comparing overlap of PBMC and TIL TCR repertoires within HNSCC patients indicates no major differences in clonal similarity based on p16 (**C**) or smoking status (**D**).

**(E)** UMAP projection of T cells containing αβ and γδ TCRs from scRNA-seq illustrates γδ T cells are predominantly found among $CD8^+$-rich phenotypes, while αβ T cells span all clusters (see also **Figure 2B**).

**(F)** Cluster composition of αβ and γδ T cells reveals that αβ T cells populate all transcriptional clusters, whereas γδ T cells are restricted to select subsets, particularly exhausted, memory-like, and bystander clusters.

**(G)** γδ T cells exhibit a distinct effector/NK-like transcriptional profile, including increased expression of *GZMB*, *ITGAM* (CD11b), *NCAM1* (CD56), and KLR molecules (*KLRC1*, *KLRD1*) compared to αβ T cells.

**(H)** Flow cytometry analysis confirms elevated CD11b (encoded by *ITGAM*) and CD56 (encoded by *NCAM1*) expression in γδ T cells relative to non-γδ T cells across PBMCs and TILs.

**(I)** Representative flow plots from patient 23 demonstrate higher abundance of γδ T cells in PBMCs than TILs and show that many γδ T cells are $CD4^-CD8^-$.

**(J)** Distribution of $CD4^+$, $CD8^+$, and double-negative subsets among all T cells (top) and γδ T cells (bottom) across HDs, PBMCs, and TILs highlights the predominance of the $CD4^-CD8^-$ phenotype in γδ cells.

**(K)** Quantification of γδ T cell frequencies reveals significantly higher levels in HD and patient-matched PBMCs compared to HNSCC TILs.

**(L)** UMAP projection of MAIT and non-MAIT T cells shows that MAIT cells are broadly distributed across most scRNA-seq clusters.

**(M)** Cluster representation analysis indicates MAIT cells are present in all clusters but at varying proportions relative to non-MAIT cells.

**(N)** Differential expression analysis identifies canonical MAIT markers (e.g., *KLRB1*) upregulated in tumor-infiltrating MAIT cells, with stemness-associated genes (e.g., *SELL, LEF1*) downregulated relative to non-MAIT cells.

**(O)** Tumor enrichment of MAIT and non-MAIT T cells. Boxplots show $\log_2$ tumor enrichment of populations; center line = median, box = interquartile range, whiskers = ±1.5×IQR.

**(P)** Flow cytometry gating of MAIT cells in PBMCs and TILs (patient 23) indicates similar frequencies between compartments.

**(Q)** Quantitative comparison of MAIT frequencies shows reduced levels in HNSCC patient samples compared to HD PBMCs, with no significant difference between HNSCC patient PBMCs and TILs.

Statistical tests: **(A-B)** mixed-effects analysis with patient-matched pairing and Fisher's LSD test; **(C-D)** unpaired t-test; **(K,Q)** one-way ANOVA with Tukey's multiple comparisons test.

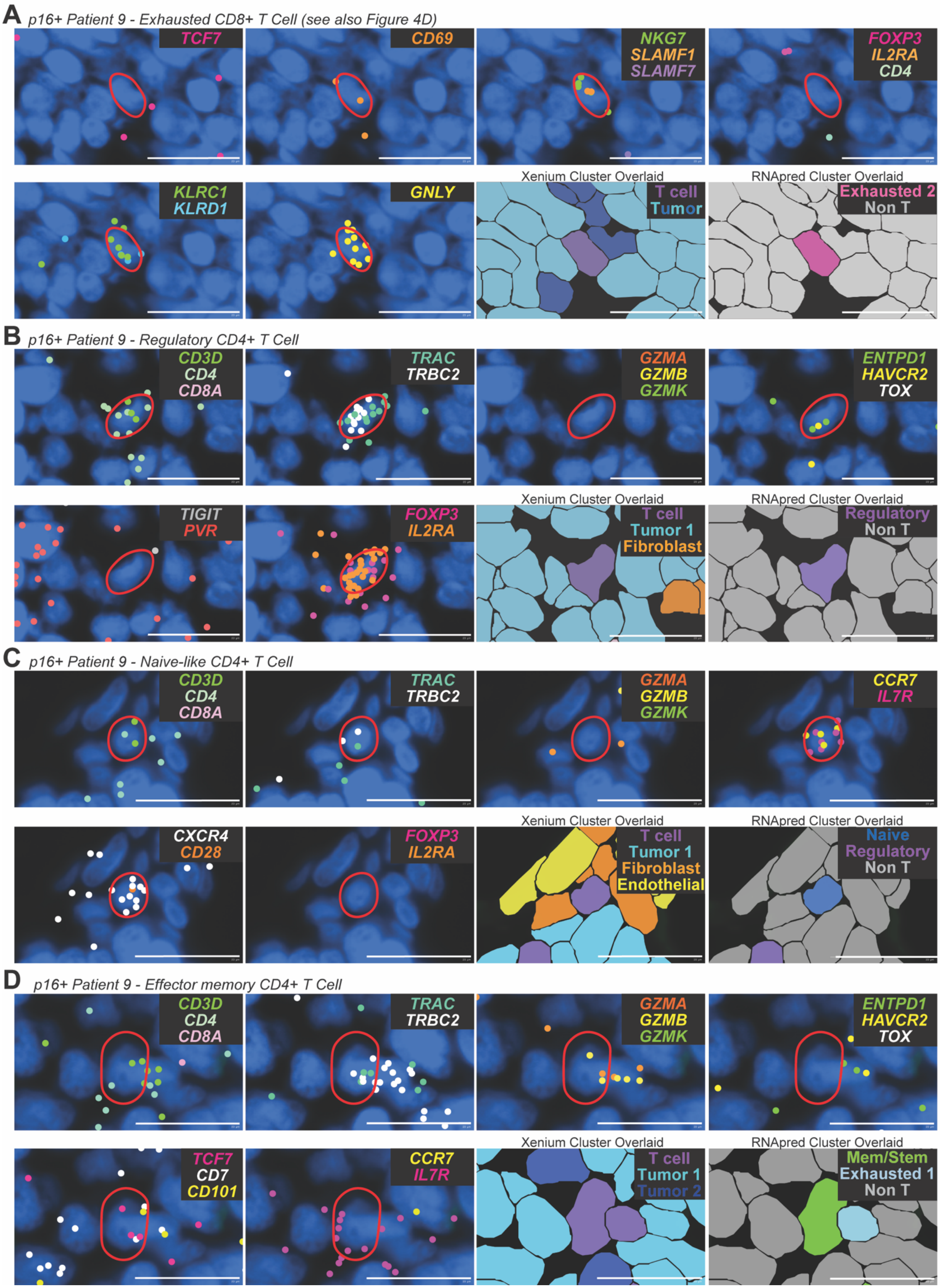
A p16+ Patient 9 - Exhausted CD8+ T Cell (see also Figure 4D)
TCF7
CD69
NKG7
SLAMF1
SLAMF7
FOXP3
IL2RA
CD4
KLRC1
KLRD1
GNLY
Xenium Cluster Overlaid
T cell
Tumor
RNApred Cluster Overlaid
Exhausted 2
Non T
B p16+ Patient 9 - Regulatory CD4+ T Cell
CD3D
CD4
CD8A
TRAC
TRBC2
GZMA
GZMB
GZMK
ENTPD1
HAVCR2
TOX
TIGIT
PVR
FOXP3
IL2RA
Xenium Cluster Overlaid
T cell
Tumor 1
Fibroblast
RNApred Cluster Overlaid
Regulatory
Non T
C p16+ Patient 9 - Naive-like CD4+ T Cell
CD3D
CD4
CD8A
TRAC
TRBC2
GZMA
GZMB
GZMK
CCR7
IL7R
CXCR4
CD28
FOXP3
IL2RA
Xenium Cluster Overlaid
T cell
Tumor 1
Fibroblast
Endothelial
RNApred Cluster Overlaid
Naive
Regulatory
Non T
D p16+ Patient 9 - Effector memory CD4+ T Cell
CD3D
CD4
CD8A
TRAC
TRBC2
GZMA
GZMB
GZMK
ENTPD1
HAVCR2
TOX
TCF7
CD7
CD101
CCR7
IL7R
Xenium Cluster Overlaid
T cell
Tumor 1
Tumor 2
RNApred Cluster Overlaid
Mem/Stem
Exhausted 1
Non T

**Figure S5. Spatial transcriptomic identification of distinct T cell phenotypes in HNSCC tumors.**

**(A)** Representative Xenium images of the exhausted $CD8^+$ T cell from patient 9 also shown in **Figure 4D**. This cell also displays robust expression of *NKG7*, *SLAMF1*, *KLRC1*, *KLRD1*, and *GNLY* transcripts. Xenium gene expression clusters overlaid onto DAPI images reveal that this T cell is surrounded by tumor cells and T cell $RNA_{pred}$ clusters overlaid onto images predicts that the cell falls within the Exhausted 2 scRNA-seq cluster based on its transcriptional profile.

**(B)** Representative Xenium images of a regulatory $CD4^+$ αβ T cell identified by co-expression of *CD3D*, *CD4*, *TRAC*, and *TRBC2*, alongside *FOXP3* and *IL2RA*, and absence of cytotoxic gene expression. Xenium gene expression cluster overlays indicate its localization within a tumor-rich region, and $RNA_{pred}$ assignment supports a regulatory phenotype.

**(C)** Representative Xenium images of a naïve-like $CD4^+$ αβ T cell characterized by expression of *CCR7*, *IL7R*, and *CXCR4*, with no detectable granzymes or regulatory markers. Overlay of the Xenium gene expression clusters show this cell positioned adjacent to fibroblasts, tumor, and endothelial cells, and $RNA_{pred}$ overlay reveals that it was assigned to the naïve/$T_{cm}$ scRNA-seq cluster.

**(D)** Representative Xenium image of a memory-like cell adjacent to an exhausted T cell. The exhausted cell expresses *GZMA, GZMB*, *ENTPD1*, and *HAVCR2*, characteristic of terminal exhaustion, while the neighboring memory-like cell expresses *TCF7* and *IL7R* without exhaustion markers. These transcriptional differences were captured by $RNA_{pred}$, which assigned $T_{em}$ and Exhausted 1 identities, respectively.

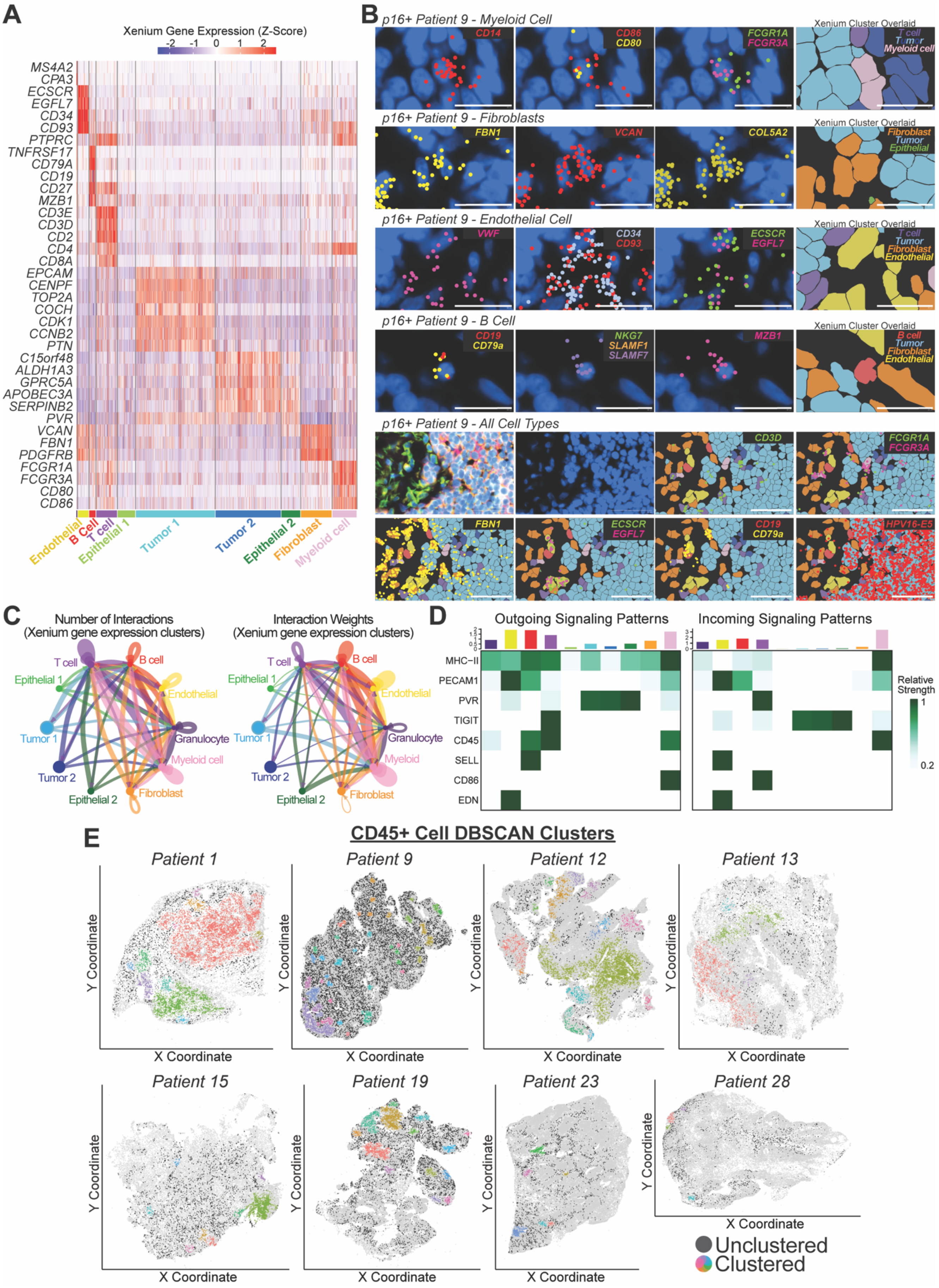

A
Xenium Gene Expression (Z-Score)
-2 -1 0 1 2
MS4A2
CPA3
ECSCR
EGFL7
CD34
CD93
PTPRC
TNFRSF17
CD79A
CD19
CD27
MZB1
CD3E
CD3D
CD2
CD4
CD8A
EPCAM
CENPF
TOP2A
COCH
CDK1
CCNB2
PTN
C15orf48
ALDH1A3
GPRC5A
APOBEC3A
SERPINB2
PVR
VCAN
FBN1
PDGFRB
FCGR1A
FCGR3A
CD80
CD86
Endothelial
B Cell
T cell
Epithelial 1
Tumor 1
Tumor 2
Epithelial 2
Fibroblast
Myeloid cell
B
p16+ Patient 9 - Myeloid Cell
CD14
CD86
CD80
FCGR1A
FCGR3A
Xenium Cluster Overlaid
T cell
Tumor
Myeloid cell
p16+ Patient 9 - Fibroblasts
FBN1
VCAN
COL5A2
Xenium Cluster Overlaid
Fibroblast
Tumor
Epithelial
p16+ Patient 9 - Endothelial Cell
VWF
CD34
CD93
ECSCR
EGFL7
Xenium Cluster Overlaid
T cell
Tumor
Fibroblast
Endothelial
p16+ Patient 9 - B Cell
CD19
CD79a
NKG7
SLAMF1
SLAMF7
MZB1
Xenium Cluster Overlaid
B cell
Tumor
Fibroblast
Endothelial
p16+ Patient 9 - All Cell Types
CD3D
FCGR1A
FCGR3A
FBN1
ECSCR
EGFL7
CD19
CD79a
HPV16-E5
C
Number of Interactions
(Xenium gene expression clusters)
Interaction Weights
(Xenium gene expression clusters)
T cell
B cell
Epithelial 1
Endothelial
Granulocyte
Tumor 1
Myeloid cell
Myeloid
Tumor 2
Fibroblast
Epithelial 2
D
Outgoing Signaling Patterns
Incoming Signaling Patterns
MHC−II
PECAM1
PVR
TIGIT
CD45
SELL
CD86
EDN
Relative Strength
1
0.2
E
CD45+ Cell DBSCAN Clusters
Patient 1
Patient 9
Patient 12
Patient 13
Patient 15
Patient 19
Patient 23
Patient 28
Y Coordinate
X Coordinate
Unclustered
Clustered

**Figure S6. Validation and analysis of Xenium-defined gene expression clusters.**

**(A)** Average expression of selected genes in Xenium-predicted gene expression clusters from all patient samples. Each column represents a single cell, grouped by annotated cluster.

**(B)** Spatial localization of lineage-defining transcripts supports cell type assignments as myeloid cells, fibroblasts, endothelial cells, and B cells. Representative high-magnification views (scale bar = 20 μm) show DAPI-stained tissue sections with individual transcripts localized to distinct cells, while overlaid colored segmentations correspond to Xenium gene expression cluster assignments. Lower-magnification images (scale bar = 50 μm) display all cell types within the same field of view, demonstrating the spatial specificity of marker gene expression within their corresponding predicted cell types.

**(C)** CellChat analysis illustrating the number of different receptor-ligand pairs (left) and strength/probability of interaction based on gene expression level of ligands and receptors (right) among Xenium gene expression clusters. Line thickness reflects interaction count or weight.

**(D)** CellChat overview of the outgoing (receptor) and incoming (ligand) signaling patterns among Xenium gene expression clusters, revealing directional communication networks within the tumor microenvironment.

**(E)** Spatial distribution of $CD45^+$ immune cell clusters identified by DBSCAN in each patient. Clusters were defined by having a minimum of 35 $CD45^+$ cells within an 80 μm radius. Colored cells represent individual DBSCAN clusters; unclustered $CD45^+$ cells are shown in black, and all remaining cells are gray.

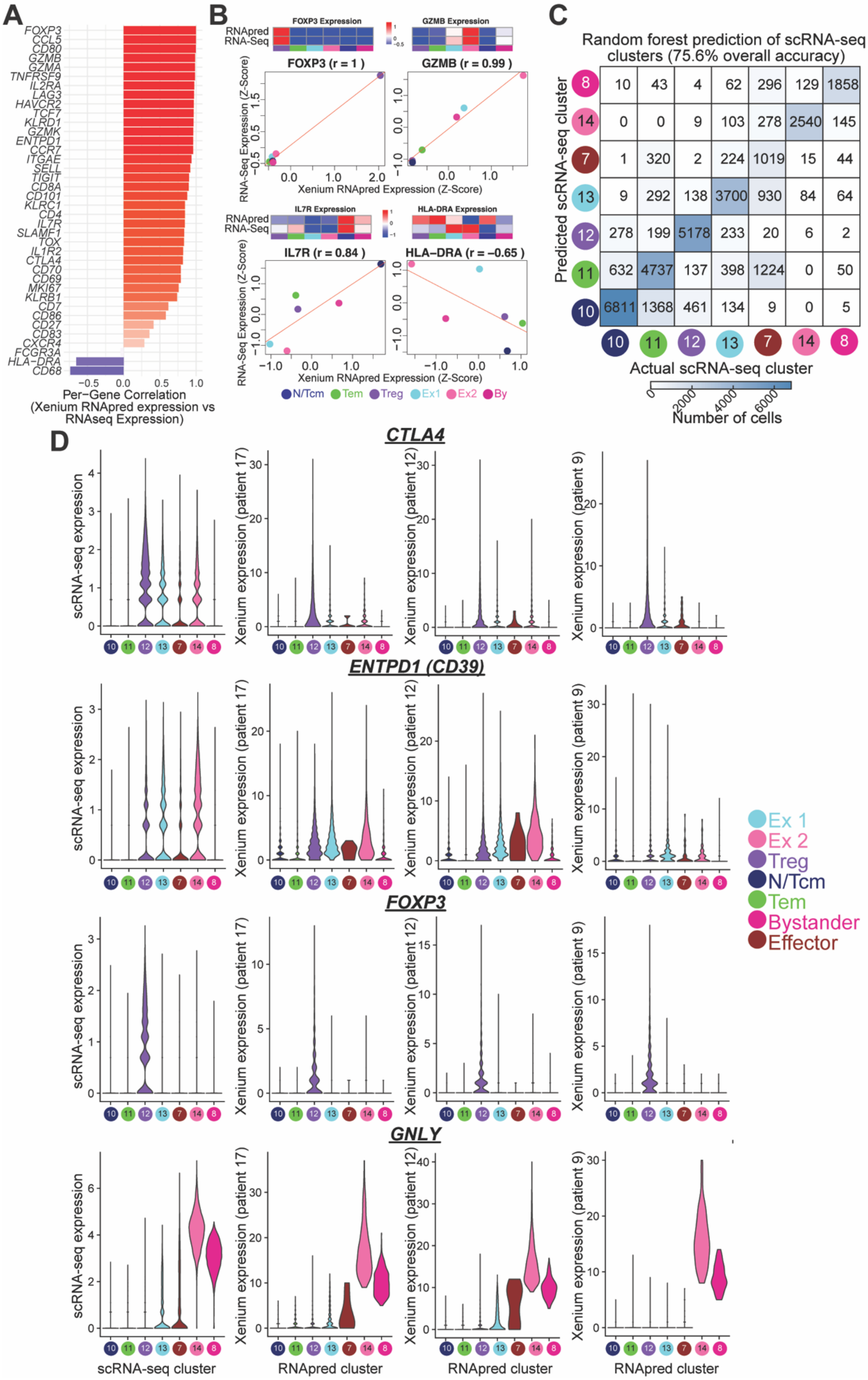

A
FOXP3
CCL5
CD80
GZMB
GZMA
TNFRSF9
IL2RA
LAG3
HAVCR2
TCF7
KLRD1
GZMK
ENTPD1
CCR7
ITGAE
SELL
TIGIT
CD8A
CD101
KLRC1
CD4
IL7R
SLAMF1
TOX
IL1R2
CTLA4
CD70
CD69
MKI67
KLRB1
CD7
CD86
CD27
CD83
CXCR4
FCGR3A
HLA−DRA
CD68
−0.5 0.0 0.5 1.0
Per-Gene Correlation
(Xenium RNApred expression vs
RNAseq Expression)
B
FOXP3 Expression
GZMB Expression
RNApred
RNA-Seq
FOXP3 (r = 1 )
GZMB (r = 0.99 )
RNA-Seq Expression (Z-Score)
Xenium RNApred Expression (Z-Score)
IL7R Expression
HLA-DRA Expression
IL7R (r = 0.84 )
HLA−DRA (r = −0.65 )
N/Tcm Tem Treg Ex1 Ex2 By
C
Random forest prediction of scRNA-seq
clusters (75.6% overall accuracy)
Predicted scRNA-seq cluster
8: 10 43 4 62 296 129 1858
14: 0 0 9 103 278 2540 145
7: 1 320 2 224 1019 15 44
13: 9 292 138 3700 930 84 64
12: 278 199 5178 233 20 6 2
11: 632 4737 137 398 1224 0 50
10: 6811 1368 461 134 9 0 5
10 11 12 13 7 14 8
Actual scRNA-seq cluster
0 2000 4000 6000
Number of cells
D
CTLA4
ENTPD1 (CD39)
FOXP3
GNLY
scRNA-seq expression
Xenium expression (patient 17)
Xenium expression (patient 12)
Xenium expression (patient 9)
10 11 12 13 7 14 8
scRNA-seq cluster
RNApred cluster
RNApred cluster
RNApred cluster
Ex 1
Ex 2
Treg
N/Tcm
Tem
Bystander
Effector

**Figure S7. Validation of scRNA-seq integration with Xenium spatial transcriptomics.**

A random forest classifier was trained using 52 genes expressed in T cells across both scRNA-seq and Xenium datasets and minimally expressed in tumor cells, minimizing segmentation or transcript misassignment errors in tumor-dense regions. The model assigned a predicted RNA-seq cluster ($RNA_{pred}$) to each Xenium T cell, excluding transient states (dividing and quiescent). The model was tested on 80% of scRNA-seq cells, with 20% held out for validation.

**(A)** Per-gene correlations between Xenium gene expression in $RNA_{pred}$ clusters and scRNA-seq gene expression in the corresponding scRNA-seq-defined clusters were calculated by z-score scaling of average expression across matched clusters within each dataset. Most genes showed strong concordance, indicating that transcriptional patterns from scRNA-seq were preserved in $RNA_{pred}$ clusters.

**(B)** Representative gene-level comparisons of gene expression between Xenium (stratified by $RNA_{pred}$) and scRNA-seq (stratified by scRNA-seq cluster). Each point represents a cluster, colored by RNA cluster identity.

**(C)** When tested on held-out scRNA-seq data, the Random forest classifier achieved 75.6% accuracy in predicting scRNA-seq cluster identity. The confusion matrix shows the number of cells in each actual scRNA-seq cluster and their predicted scRNA-seq cluster.

**(D)** scRNA-seq-based expression of selected genes in scRNA-seq clusters (left) and expression of these genes in Xenium data, stratified by $RNA_{pred}$ cluster. Three representative patients are shown; these patients are highlighted in Figures 6-8.

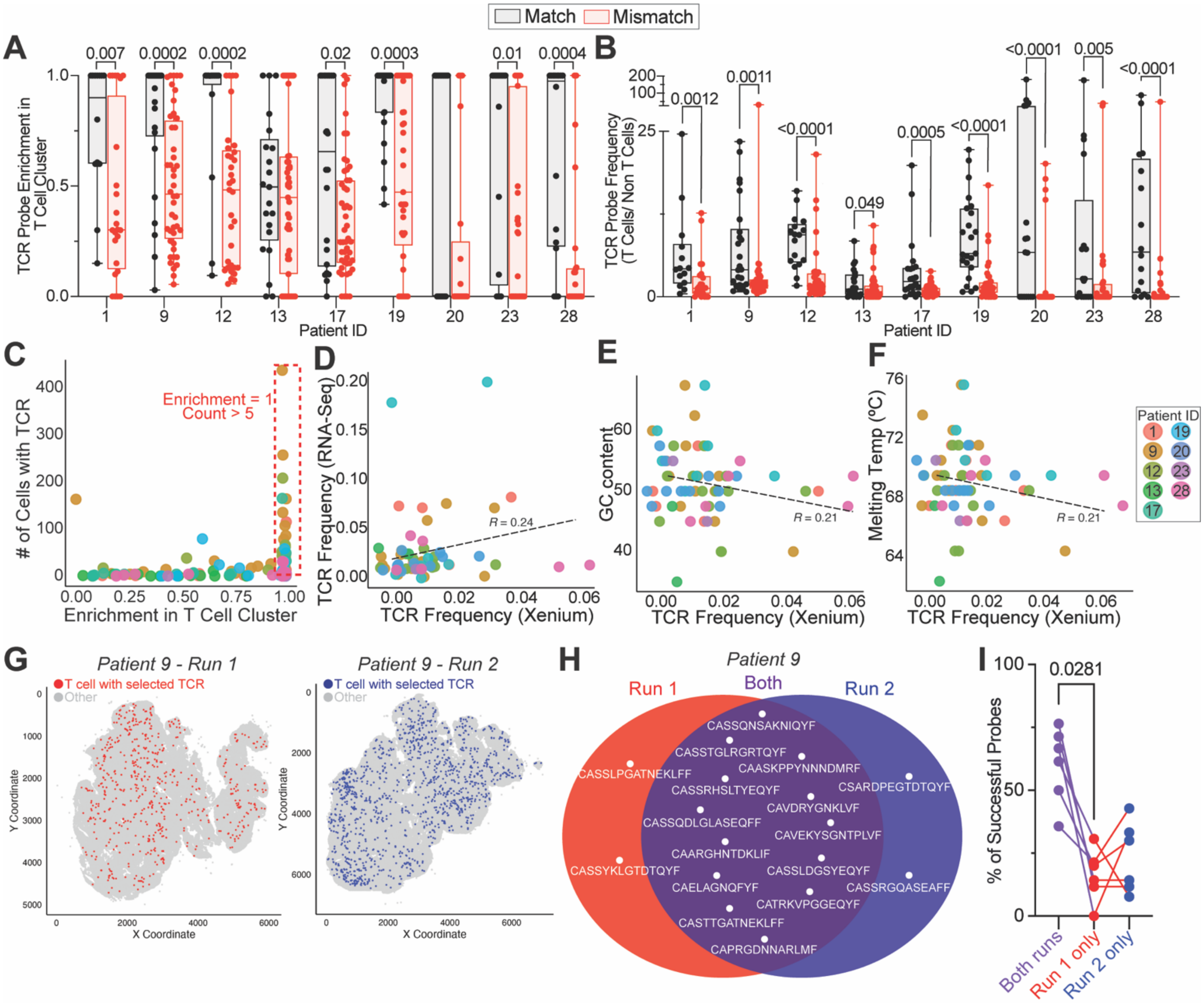
Match
Mismatch
A
TCR Probe Enrichment in T Cell Cluster
0.007
0.0002
0.0002
0.02
0.0003
0.01
0.0004
Patient ID
B
TCR Probe Frequency (T Cells/ Non T Cells)
0.0012
0.0011
<0.0001
0.049
0.0005
<0.0001
<0.0001
0.005
<0.0001
Patient ID
C
### of Cells with TCR
Enrichment = 1
Count > 5
Enrichment in T Cell Cluster
D
TCR Frequency (RNA-Seq)
R = 0.24
TCR Frequency (Xenium)
E
GC content
R = 0.21
TCR Frequency (Xenium)
F
Melting Temp (ºC)
R = 0.21
TCR Frequency (Xenium)
Patient ID
G
Patient 9 - Run 1
T cell with selected TCR
Other
Y Coordinate
X Coordinate
Patient 9 - Run 2
T cell with selected TCR
Other
Y Coordinate
X Coordinate
H
Patient 9
Run 1
Both
Run 2
CASSQNSAKNIQYF
CASSTGLRGRTQYF
CAASKPPYNNNDMRF
CASSLPGATNEKLFF
CASSRHSLTYEQYF
CSARDPEGTDTQYF
CAVDRYGNKLVF
CASSQDLGLASEQFF
CAVEKYSGNTPLVF
CAARGHNTDKLIF
CASSYKLGTDTQYF
CASSLDGSYEQYF
CAELAGNQFYF
CASSRGQASEAFF
CATRKVPGGEQYF
CASTTGATNEKLFF
CAPRGDNNARLMF
I
0.0281
% of Successful Probes
Both runs
Run 1 only
Run 2 only

**Figure S8. Validation of custom TCR-targeting Xenium probes.**

**(A)** Enrichment of patient-matched versus mismatched TCR probes within the Xenium-defined T cell gene expression cluster reveals that matched probes are significantly more enriched in the T cell cluster than mismatched probes. Matched probes were designed using TCR sequences derived from the same patient in which the tissue was profiled, ensuring both probe and tissue originated from the same individual. Mismatched probes target TCRs from a different patient than the one whose tissue was profiled. For each probe, TCR frequency across gene expression clusters was normalized to the maximum value for that probe, yielding relative enrichment values from 0 to 1. Each point represents one TCR probe. P values were calculated using Mann-Whitney tests, using the FDR p-value correction for multiple corrections.

**(B)** Frequency of matched and mismatched TCR probes in T cells versus non T cells shows that patient-matched probes are preferentially detected in T cells. P values were calculated as in **(A)**.

**(C)** Number of cells with a TCR detected versus enrichment in the T cell cluster. Red box denotes the selection criteria used for downstream analysis: count > 5 and relative enrichment = 1. A total of 61 TCR probes expressed in over 2,000 T cells in eight patients met these criteria.

**(D-F)** Comparison of Xenium-detected TCR frequencies with scRNA-seq TCR frequencies (**D**), probe GC content (**E**), and melting temperature (**F**) across successful probes.

**(G)** Spatial distribution of T cells with selected TCRs found in patient 9 tissue during Xenium run one (left) versus run two (right). In both sections, TCRs are dispersed throughout the whole tissue.

**(H)** Comparison of patient 9 TCR probes across technical replicate tissue sections reveals that most probes were consistently detected, though two were unique to each run.

**(I)** Percentage of matched TCR probes successfully detected in both technical replicates (purple), only in run one (red), or only in run two (blue) across patient samples. Each line represents one patient. P value calculated using Friedman test.

**Table S1. Overview of HNSCC patient counts and clinical characteristics.** Summary of clinical features for all patients included in the study. p16 status was assessed only for oropharyngeal tumors; all others were classified as p16-negative for this study. Pack-years were calculated as the number of cigarette packs smoked per day multiplied by the number of years the individual has smoked.

| Group | N |
|---|---|
| **Total Counts** | |
| TILs from HNSCC patient | 27 |
| PBMCs from HNSCC patient | 26 |
| PBMCs from healthy donor | 5 |
| Tumors examined on Xenium | 10 |
| **HNSCC patient p16 Status** | |
| p16- | 20 |
| p16+ | 7 |
| **HNSCC patient Smoking Status** | |
| Pack-years = 0 | 11 |
| Pack-years >0 | 16 |
| **Tumor Site** | |
| Oropharynx (OP) | 8 |
| Oral cavity (OC) | 10 |
| Hypopharynx (HP) | 3 |
| Larynx (LA) | 6 |
| **Tumor Recurrence** | |
| Primary | 23 |
| Recurrent | 4 |

**Table S2. Per-patient clinical metadata for the HNSCC cohort.** "NA" indicates not tested for p16 status. Smoking is measured in pack years. Tumor sites include oropharynx (OP), oral cavity (OC), hypopharynx (HP), and larynx (LA). Previous treatment includes chemoradiation (CXRT) and radiation (XRT). Alternate IDs are used in some deposited Xenium files.

| Patient ID | Primary vs recurrent | p16 status | Smoking pack years | Tumor site | Subsite | Previous treatment | Tumor | Nodes | Metastases | Alternate ID |
|---|---|---|---|---|---|---|---|---|---|---|
| 1 | recurrent | positive | 50 | OP | tonsil | CXRT | 1 | 0 | 0 | 230322 |
| 2 | primary | NA | 0 | OC | tongue | none | 2 | 0 | 0 | 230410 |
| 3 | primary | NA | 100 | HP | NA | none | 2 | 3 | 0 | 230424 |
| 4 | primary | NA | 50 | OC | tongue | none | 3 | 3b | 0 | 230517 |
| 5 | primary | NA | 40 | LA | glottis | none | 4 | 0 | 0 | 230717 |
| 6 | primary | NA | 0 | LA | glottis | none | 4a | 3b | 0 | 230724 |
| 7 | primary | NA | 0 | LA | multiple | none | 4a | 3b | 0 | 230809 |
| 8 | primary | NA | 65 | OC | tongue | none | 3 | 2b | 0 | 230816 |
| 9 | primary | positive | 10 | OP | base of tongue | none | 3 | 2b | 0 | 230913 |
| 10 | | | | | ameloblastoma | | | | | 230925 |
| 11 | primary | NA | 40 | LA | glottis | none | 2 | 0 | 0 | 231004A |
| 12 | primary | positive | 44 | OP | NA | none | 4 | 2c | 0 | 231004B |
| 13 | recurrent | positive | 0 | OP | tonsil | CXRT | 4 | a | 0 | 231101 |
| 14 | primary | NA | 20 | OC | tongue | none | 3 | 0 | 0 | 231108 |
| 15 | primary | NA | 0 | OC | alveolus | none | 4a | 3b | 1 | 231110 |
| 16 | primary | NA | 25 | LA | glottis | none | 4a | 2b | 0 | 231113 |
| 17 | primary | NA | 40 | OC | floor of mouth | none | 4a | 0 | 0 | 231120 |
| 18 | primary | positive | 0 | OP | base of tongue | none | 4 | 0 | 0 | 231220 |
| 19 | primary | NA | 80 | LA | supraglottis | none | 3 | 3b | 0 | 240103 |
| 20 | primary | negative | 42 | HP | pyriform | XRT | 3 | 3b | 0 | 240109 |
| 21 | primary | positive | 40 | OP | soft palate | none | 2 | 1 | 0 | 240207 |
| 22 | recurrent | negative | 0 | OC | gingiva | CXRT | 4 | 0 | 0 | 240212 |
| 23 | primary | NA | 45 | OC | tongue | none | 2 | 0 | 0 | 240228 |
| 24 | primary | NA | 0 | OC | tongue | none | 4 | 3b | 0 | 240322 |
| 25 | primary | negative | 30 | OP | tonsil/base of tongue | none | 4 | 2c | 0 | 240326 |
| 26 | primary | NA | 0 | OC | tongue | none | 4 | 3b | 0 | 240402 |
| 27 | primary | negative | 30 | HP | pharyngeal wall | none | 4 | 3b | 0 | 240425 |
| 28 | recurrent | positive | 0 | OP | tongue | CXRT | 2 | 0 | 0 | 240501 |

**Table S3. Human antibodies used in spectral cell sort.**

| Antibody | Source | Catalogue # | Clone | µl per 50µL |
|---|---|---|---|---|
| TCR v$\alpha$7.2 Biotin | BioLegend | 351724 | 3C10 | 1 |
| Ghost UV450 | Tonbo | 13-0868-T100 | - | 1 |
| CD16 BUV737 | ThermoFisher | 367-0168-41 | CB16 | 0.25 |
| Human TruStain FcX | BioLegend | 422302 | - | 0.5 |
| CD69 BUV395 | ThermoFisher | 363-0699-41 | FN50 | 1 |
| CD14 BUV496 | ThermoFisher | 364-0149-41 | 61D3 | 1 |
| CD45 BUV563 | ThermoFisher | 365-0459-41 | H130 | 1 |
| CD19 BUV615 | ThermoFisher | 366-0198-42 | SJ25C1 | 1 |
| LAG3 BUV805 | ThermoFisher | 368-2239-41 | 3DS223H | 1 |
| TIM3 BV421 | BioLegend | 345008 | F38-2E2 | 1 |
| HLA-DR Pacific Blue | BioLegend | 260164 | L243 | 0.25 |
| CD8 BV510 | BioLegend | 344731 | SK-1 | 0.5 |
| CD45RA BV605 | BioLegend | 304134 | HI100 | 1 |
| CD101 BV650 | BD | 747547 | V7.1 | 1 |
| CD56 BV711 | BioLegend | 362541 | 5.1H11 | 1 |
| CD127 BV750 | BD | 747089 | HIL-7R-M21 | 1 |
| PD-1 BV786 | BioLegend | 329929 | EH12.2H7 | 1 |
| CD3 AF488 | BioLegend | 300320 | UCHT1 | 1 |
| Streptavidin Spark Blue 574 | BioLegend | 405355 | - | 0.25 |
| IgD PerCP-Cy5.5 | BioLegend | 348208 | 1A6-2 | 1 |
| CD11b PerCP-eFluor-710 | ThermoFisher | 46-0112-82 | M1/70 | 1 |
| CD27 RB744 | BD | 570713 | M-T271 | 0.125 |
| CD39 PerCP-Fire806 | BioLegend | 328249 | A1 | 1 |
| CD28 PE | BioLegend | 302907 | CD28.2 | 1 |
| TCRγδ PE-Vio615 | Miltenyi Biotec | 130-131-974 | REA591 | 1 |
| TIGIT PE-Cy5 | BioLegend | 372747 | A15153G | 1 |
| CD38 PE-Fire 700 | BioLegend | 397121 | S17015A | 0.5 |
| CD7 PE-Cy7 | BioLegend | 343113 | CD7-6B7 | 0.25 |
| CCR7 Spark NIR 685 | BioLegend | 353258 | G043H7 | 1 |
| CD11c APC-Fire 750 | BioLegend | 337239 | Bu15 | 1 |
| CD4 APC-Fire 810 | BioLegend | 344661 | SK-3 | 0.25 |
| Siglec-7 APC | BioLegend | 347705 | S7.7 | 1 |
| TotalSeq™-C0251 anti-human Hashtag 1 | BioLegend | 394661 | 2M2 | 1 |
| TotalSeq™-C0252 anti-human Hashtag 2 | BioLegend | 394663 | 2M2 | 1 |
| TotalSeq™-C0253 anti-human Hashtag 3 | BioLegend | 394665 | 2M2 | 1 |
| TotalSeq™-C0254 anti-human Hashtag 4 | BioLegend | 394667 | 2M2 | 1 |

**Table S5. Custom add-on panel employed in Xenium spatial transcriptomics to target key T cell and tumor cell genes and HPV16 transcripts.**

| Gene | Ensembl ID | Probes |
|---|---|---|
| *NFE2L2* | ENSG00000116044 | 8 |
| *KEAP1* | ENSG00000079999 | 8 |
| *TXNRD1* | ENSG00000198431 | 8 |
| *AKR1C3* | ENSG00000196139 | 8 |
| *AICDA* | ENSG00000111732 | 8 |
| *BCL6* | ENSG00000113916 | 8 |
| *IGHD* | ENSG00000211898 | 8 |
| *PIGR* | ENSG00000162896 | 8 |
| *CD7* | ENSG00000173762 | 8 |
| *CD101* | ENSG00000134256 | 8 |
| *TIGIT* | ENSG00000181847 | 8 |
| *TOX* | ENSG00000198846 | 8 |
| *TCF7* | ENSG00000081059 | 8 |
| *HLA-DRA* | ENSG00000204287 | 5 |
| *ENTPD1* | ENSG00000138185 | 8 |
| *ITGAE* | ENSG00000083457 | 8 |
| *PVR* | ENSG00000073008 | 8 |
| *CD226* | ENSG00000150637 | 8 |
| *CD80* | ENSG00000121594 | 8 |
| *TRDC* | ENSG00000211829 | 8 |
| *TRBC1* | ENSG00000211751 | 8 |
| *TRBC2* | ENSG00000211772 | 8 |
| *HPV16-E1* | HPV16-E1 | 5 |
| *HPV16-E2* | HPV16-E2 | 2 |
| *HPV16-E4* | HPV16-E4 | 1 |
| *HPV16-E5* | HPV16-E5 | 2 |
| *HPV16-E6* | HPV16-E6 | 2 |
| *HPV16-E7* | HPV16-E7 | 2 |
| *HPV16-L1* | HPV16-L1 | 7 |
| *HPV16-L2* | HPV16-L2 | 4 |
